# Lipid droplets sequester cell death effectors and delay regulated cell death execution

**DOI:** 10.64898/2026.03.16.712084

**Authors:** Yingquan Shan, Kinga B Stopa-Kerkhofs, Rouchidane Eyitayo Akandé, Florence Jollivet, Victor Girard, Catherine Jamard, Lena Sapozhnikov, Eli Arama, Judit Szécsi, Mohammed Bendahmane, Nathalie Davoust, Ludivine Walter, Mingyao Liu, Nicolas Aznar, Gabriel Ichim, Bertrand Mollereau

**Author notes:** Correspondence (ARE); (G.I.); (B.M.). These authors contributed equally to this work (Yingquan Shan, Kinga B Stopa-Kerkhofs, Rouchidane Eyitayo Akandé).

## Abstract

Normal and cancer cells accumulate lipid droplets (LDs) under stress to buffer lipotoxicity, but their role in regulated cell death (RCD) remains incompletely understood. Here, we explored LD accumulation across multiple apoptotic and non-apoptotic RCD modalities in human cancer cells and *Drosophila* germ cells. We found that LD accumulation arises from *de novo* LD biogenesis, whereas LD lipolysis remains active—or even enhanced—in dying germ cells and cancer cells, respectively. In *Drosophila*, LD accumulation in ATGL/Bmm lipase loss of function mutant inhibited germ cell death, supporting a protective role of LDs. Proteomic and imaging analyses revealed a broad redistribution of LD-associated proteins, encompassing lipid metabolism and stress response factors, as well as the pro-apoptotic effector Bax in human cancer cells during cell death. Enhanced LD–mitochondria contacts promoted the translocation of conformationally active Bax from mitochondria to LDs, thereby delaying apoptosis execution. Consistently, LD depletion sensitizes cells to Bax-dependent apoptosis, whereas LD accumulation confers resistance. Collectively, these findings define LD accumulation during cell death as a delaying mechanism in which LDs sequester mitochondrial cell death regulators, attenuating their pro-death activity and revealing potential therapeutic implications for apoptosis-resistant cancers.

## Introduction

Lipids are essential for a wide array of cellular functions, including membrane biogenesis, energy metabolism, extracellular and intercellular signaling, and the regulation of cell death. Within cells, fatty acids (FAs)—whether acquired exogenously or synthesized *de novo*—are stored in lipid droplets (LDs), specialized organelles composed of a neutral lipid core, primarily triacylglycerols (TGs) and sterol esters (SEs), surrounded by a phospholipid monolayer enriched with various LD-associated proteins ^1^. LD homeostasis has emerged as a critical process in tissue physiology, particularly in adipose tissue and the central nervous system. Disruption of LD dynamics has been implicated in numerous pathological conditions, including obesity, atherosclerosis, hepatic steatosis, and neurodegenerative diseases ^2, 3^. Notably, aberrant LD accumulation is also recognized as a hallmark of resistance to cancer treatments, although the underlying molecular mechanisms remain unclear ^4–6^.

The regulation of lipid storage and mobilization within LDs relies on a network of evolutionarily conserved anabolic and catabolic enzymes. *Drosophila melanogaster* has been a pioneering model in LD research, providing key insights into both canonical and non-canonical LD functions ^7, 8^. LD biogenesis initiates at the endoplasmic reticulum (ER) membrane, where enzymes such as diacylglycerol acyltransferase 1 (DGAT1), Midway (Mdy) in *Drosophila* and DGAT2 catalyze the rate-limiting step of TG synthesis. Conversely, lipolysis is mediated by enzymes such as adipose triglyceride lipase (ATGL), known as Brummer (Bmm) in *Drosophila* ^9^. The activity of lipases is modulated by perilipins (PLINs), a family of LD-associated proteins that stabilize LDs, limit basal lipolysis, and facilitate interactions with mitochondria ^10^. In humans, five PLIN proteins (PLIN1–5) are known ^11^, while *Drosophila* expresses two perilipins—*Lsd-1* and *Lsd-2* (dPlin1 and dPlin2) with distinct functions ^7^.

Beyond their classical role in lipid storage, LDs are increasingly recognized for their involvement in diverse cellular processes, particularly in stress responses ^8–12^. Studies in *Drosophila* and vertebrate model organisms have shown that glial cells accumulate LDs to mitigate oxidative stress and lipid peroxidation, both during development and in adulthood ^13–16^. In cancer cells, LDs similarly act as antioxidant organelles that limit lipid peroxidation and confer resistance to ferroptosis ^17^.

In addition to this established antioxidant function preventing ferroptosis execution, emerging evidence reveals further regulatory roles for LDs in controlling cell death execution. LD metabolism appears tightly interconnected with apoptosis, suggesting a bidirectional interplay. For instance, mitochondrial dysfunction triggered by intrinsic apoptotic stimuli can promote LD formation ^18, 19^. Conversely, LD-associated proteins such as members of the Bcl-2 family can modulate apoptotic signaling. Notably, Bax and Bcl-xL contain a motif enabling their binding to LDs ^20^. These observations point to a broader regulatory role for LD-associated proteins in shaping cell death outcomes, although the precise mechanisms and the contribution of distinct death modalities—such as apoptosis or necrosis—remain incompletely defined.

LDs are also detected during *Drosophila* spermatogenesis, when a significant fraction of germ cells undergo programmed death ^21–24^, further underscoring the potential involvement of LDs in the broader regulation of cell death processes across different species and physiological contexts.

In this study, we examined LD accumulation across multiple apoptotic and non-apoptotic regulated cell death modalities in human cancer cells and *Drosophila* germ cells. We show that LDs are formed through *de novo* biogenesis despite active or even elevated lipolysis. In *Drosophila*, loss of the Bmm lipase suppresses germ cell death, supporting a cytoprotective role for LDs. In human cancer cells, quantitative proteomics uncovered an extensive remodeling of the LD proteome, with the recruitment of lipid metabolism and stress response proteins. Targeted western blot analyses revealed the accumulation of the pro-apoptotic effector Bax at LDs. Enhanced LD–mitochondria contacts promoted Bax translocation from mitochondria to LDs, delaying apoptosis. Furthermore, LD depletion sensitizes cells to Bax-dependent apoptosis, whereas LD accumulation confers resistance, establishing LD accumulation as a conserved mechanism that transiently restrains mitochondrial death effectors.

## Results

### LDs accumulate during multiple RCD modalities in both human cancer cells and *Drosophila* germ cells

Normal and cancer cells frequently accumulate LDs under various stresses, including apoptosis, yet this phenomenon has not been systematically investigated using different RCD stimuli ^8, 18^. We thus examined LD accumulation with the neutral lipid dye BODIPY^493/503^ or LipidTOX, that labels neutral lipids including TGs and SEs using different apoptotic and non-apoptotic stimuli in both cancer cells and *Drosophila* models. For cancer cells, we first chose pancreatic cell line BxPC3 that is competent to intrinsic and extrinsic apoptosis, necrosis and necroptosis, while colorectal cell line HCT116 is competent to all these cell death forms except necroptosis (**Fig. S1A-I**). For intrinsic apoptosis triggered by combination of actinomycin D (ActD) and the BH3 mimetic ABT-737 (ActD/ABT-737), which induces Bax-dependent apoptosis in HCT116 cells ^25^, caspase activity was inhibited with the pan-caspase inhibitor Q-VD-OPh (QVD) to prevent cell detachment and thus enable LD detection within 18 h of cell death induction. We observed LD accumulation in both size and number during apoptosis and necrosis, with the exception of necroptosis (**Fig. 1A-D**). In general, the accumulation of LDs (size and total LD area per cell) was more robust in apoptosis compared to necrosis. To test the generality of these findings, we examined *Drosophila* male germ cells, an *in vivo* model where LDs play a key physiological role in germ cell homeostasis ^21^. Notably, this physiological germ cell death represents the sole mitochondrial-dependent cell death pathway in *Drosophila* in which LDs are readily detectable. During the four rounds of mitosis, approximately 20% of germ cells undergo spontaneous RCD characterized by loss of the plasma and nuclear membrane permeability—hallmarks of necrosis (**Fig. S2A**) ^22^. However, this is not necroptosis because *Drosophila* lacks the effectors RIPK and MLKL proteins ^26^. This germ cell death proceeds independently of active caspases while relying on mitochondrial factors, including Bcl-2 family homologs Debcl and Buffy, as well as lysosomal components ^21, 22^. Using the germ cell driver *nanos (nos-Gal4)* driving the expression of nuclear proteins Lamin-GFP and RFP-nls, we could observe that the light-refractive dying germ cell cysts (hereafter “dying germ cells”) identified by brightfield microscopy exhibit the loss of Lamin nuclear membrane localization and RFP-nls nuclei localization, which are signs of nuclear membrane permeability (**Fig. S2B**). LDs (BODIPY^+^) were observed in healthy germ cells but increased in both size and number in dying germ cells labeled with propidium iodide or LysoTracker dyes in the apical part of the testes, where germ cell death usually occurs (**Fig. 1E-G**). In addition, LD accumulation during cell death was exclusively observed in dying germ cells labeled with Vasa-EGFP, but not in somatic cyst cells expressing membrane-targeted GFP (mCD8-GFP) under the *C587-Gal4* driver **(Fig. S2C-D).** Interestingly, the accumulation of LDs was not only observed in germ cells during physiological cell death but also when apoptosis was induced by overexpressing a constitutively active Drice transgene (*actDrice*) under the *nos* driver (**Fig. S2E-G**). Collectively, the accumulation of LDs seems to be a universal response of both human cancer cells and *Drosophila* germ cells to cell death stimuli.

**Fig. 1.**
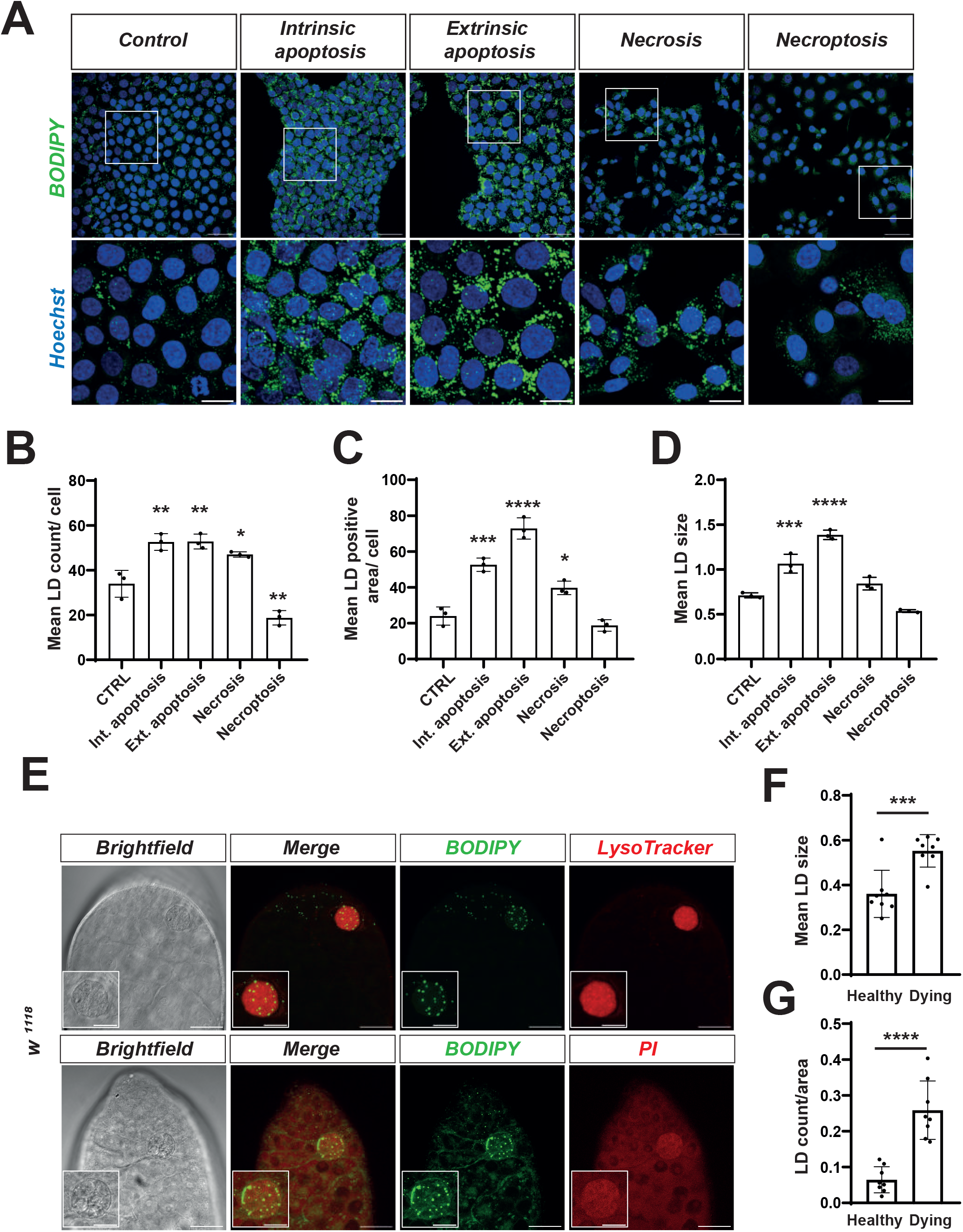
Cell death–specific accumulation of lipid droplets in cancer and *Drosophila* germ cells. **(A)** Representative fluorescence images of LDs in BxPC3 cells 18 h after induction of cell death. Cells were stained with BODIPY^493/503^ (green) to visualize LDs, and Hoechst 33342 (blue) to label nuclei. Treatments included: intrinsic apoptosis induced by Actinomycin D (ActD, 1 µM) + ABT-737 (1 µM), extrinsic apoptosis induced by cycloheximide (CHX, 5 µg/mL) + tumor necrosis factor alpha (TNFα, 20 ng/mL), necrosis induced by ionomycin (25 µM), and necroptosis by the SMAC mimetics (BV6,10 µM) + TNFα + caspase inhibitor ZVAD (10 µM). Caspase inhibitor (QVD,10 µM) was used to prevent cell detachment during apoptosis. Solid white squares indicate regions of interest, which are shown at higher magnification in the bottom panels. Scale bars: 50 µm. Scale bars in magnified images: 20 µm. **(B)** Quantification of the mean LDs count per cell. **(C)** Total LD area per cell (µm^2^). **(D)** Mean size of individual LD area (µm^2^). Data in **(B–D)** represent the mean ± SD from three independent experiments (N = 3). Statistical analysis was performed using one-way ANOVA. *, p < 0.05; **, p < 0.01; ***, p < 0.001; ****, p < 0.0001 versus untreated control (NT). **(E)** Representative images of the apical tip of testes from control (*w¹¹¹⁸*) flies. Dying germ cells are observed in differential interference contrast (Brightfield) images. The testes were stained with BODIPY^493/503^ combined with LysoTracker (upper panel) or propidium iodide (PI, lower panel). Scale bars: 20 μm. White squares at bottom left indicate magnified regions of dying germ cells. Scale bars in magnified images: 10 μm. **(F, G)** Quantification of LD size and number in healthy versus dying germ cells (n = 8/group). Statistical analysis was performed using two-tailed unpaired Student’s t-test: ***, p < 0.001; ****, p < 0.0001.

### Conserved mechanisms control LD turnover during cell death in human cancer cells and *Drosophila*

LD homeostasis depends on the balance between *de novo* LD synthesis and lipolysis ^1^. While enhanced LD formation through the *de novo* pathway has been proposed as a mechanism underlying LD accumulation during intrinsic apoptosis ^18^, the dynamics of LD turnover and the behavior of perilipin-defined LD subpopulations remain poorly characterized. Perilipins are key regulators of LD stability and accessibility to lipases ^27^. We focused on PLIN1, PLIN2, and PLIN3, the primary perilipins expressed in BxPC3 and HCT116 cancer cells (**Fig. S3A**). Of these, only PLIN2 and PLIN3 localized to LDs labelled with BODIPY **(Fig. 2A)**. Although total protein levels of PLIN2 and PLIN3 remained unchanged following apoptosis induction **(Fig. S3B, C)**, BODIPY-based immunofluorescence revealed a marked accumulation of PLIN3, but not PLIN2, on BODIPY-positive LDs after 18 h of apoptotic treatment (**Fig. 2A, S3D**). To extend our observations, we performed LD isolation that allows quantification of LD proteins by western blot. For this we used HCT116 cells, which are apoptosis-competent and express higher levels of ATGL compared to BxPC3 cells (**Fig. S1F-I, S3E**). HCT116 cells also provided sample material for LD isolation due to their robust growth characteristics. The specificity of the isolation protocol was confirmed by western blotting, which showed that LD-containing fractions (1–4) were enriched in PLIN2 and free of contamination from cytosolic (GAPDH) or mitochondrial (COX IV) proteins (**Fig. S3F**). This approach also allowed us to examine LD accumulation during the early phase of apoptosis, revealing an early rise in LD abundance detectable by BODIPY staining as soon as 5 h after induction of intrinsic apoptosis with ActD/ABT-737 (**Fig. S3G**). Consistent with PLIN3 enrichment at LDs in BxPC3 cells (**Fig. 2A, S3D**), LD accumulation in HCT116 cells was also associated with an increase in PLIN3, but not PLIN2 or ATGL within isolated LD fractions 5 hours after ActD/ABT-737 treatment (**Fig. 2B–F**). Given that PLIN3 has been previously shown to label nascent LDs ^28^, these findings, together with PLIN3 immunostaining (**Fig. 2A**), suggest that *de novo* LD biogenesis contributes to LD accumulation during intrinsic apoptosis. Supporting this interpretation, co-treatment with DGAT1/2 inhibitors and ActD/ABT-737 abolished *de novo* LD formation, as indicated by the absence of BODIPY-labeled LDs in HCT116 cells after 18 h of ActD/ABT-737 treatment (**Fig. S4A–C**), while a similar effect was observed at early stage of apoptosis (5 h of pro-apoptotic treatment) (**Fig. S4D-H)**. In these conditions, DGAT inhibition not only prevented the apoptosis-induced increase in PLIN3 but also reduced PLIN2 and ATGL levels in isolated LD fractions, providing additional evidence that *de novo* LD biogenesis is activated during intrinsic apoptosis (**Fig. S4D–H**). Moreover, the selective accumulation of PLIN3- but not PLIN2-positive LDs during apoptosis suggest the coexistence of distinct LD subpopulations, with PLIN2-positive LDs likely undergoing faster lipolysis than their PLIN3-positive counterparts. Consistent with this, PLIN3 levels increased in the LD fraction during apoptosis, whereas ATGL did not display a corresponding rise (**Fig. 2B, F**), supporting reduced lipolysis in PLIN3-marked LDs. This interpretation is further reinforced by the observation that PLIN2- but not PLIN3-positive LDs decrease when apoptosis occurs in the presence of DGAT inhibitors (**Fig. S4F, G**). To verify that PLIN2 LD subpopulation undergo efficient lipolysis, ATGL activity was inhibited during apoptosis induction. For that, we evaluated PLIN2 and ATGL levels on LD fractions of HCT116 cells in the presence of ATGL inhibitor. Inhibition of ATGL led to an increase of PLIN2 and ATGL on LD fractions (**Fig. 2G-K**). The rise in ATGL reflects the accumulation of the enzyme in its inactive, LD-bound form following lipolysis blockade (**Fig. 2K**). The increase in PLIN2 suggests that normally digested PLIN2-positive LDs become stabilized when ATGL is inhibited (**Fig. 2J**). This is consistent with active lipolysis of the PLIN2-positive LD subpopulation during apoptosis, which is prevented upon ATGL inhibition. To directly investigate LD lipolysis during apoptosis, we performed a pulse-chase assay using BODIPY ^558^/^568^ C12–labeled fatty acids, a fluorescent fatty acid analog that accumulates in LDs and enables tracking of LD-derived fatty acid mobilization over time ^29^. Upon induction of apoptosis, BODIPY ^558^/^568^ C12 fluorescence, which initially localized to LDs co-labeled with BODIPY ^493/503^, progressively decreased in ActD/ABT-737–treated HCT116 cells (**Fig. 2L, M**). To rule out a potential bias in the Red C12/LD ratio caused by LD accumulation during apoptosis, we quantified the amount of free BODIPY ^558^/^568^ C12 in Vinculin-labelled cytosolic fractions that are depleted of LDs (**Fig. 2N**). We observed a marked increase in cytosolic BODIPY ^558^/^568^ C12 following apoptosis induction in HCT116 cells (**Fig. 2O**), providing further evidence that intrinsic apoptosis is also accompanied by enhanced lipolysis. Notably, this elevated lipolytic activity indicates that apoptosis-associated LD accumulation results primarily from increased *de novo* lipid synthesis rather than from reduced LD breakdown.

**Fig. 2.**
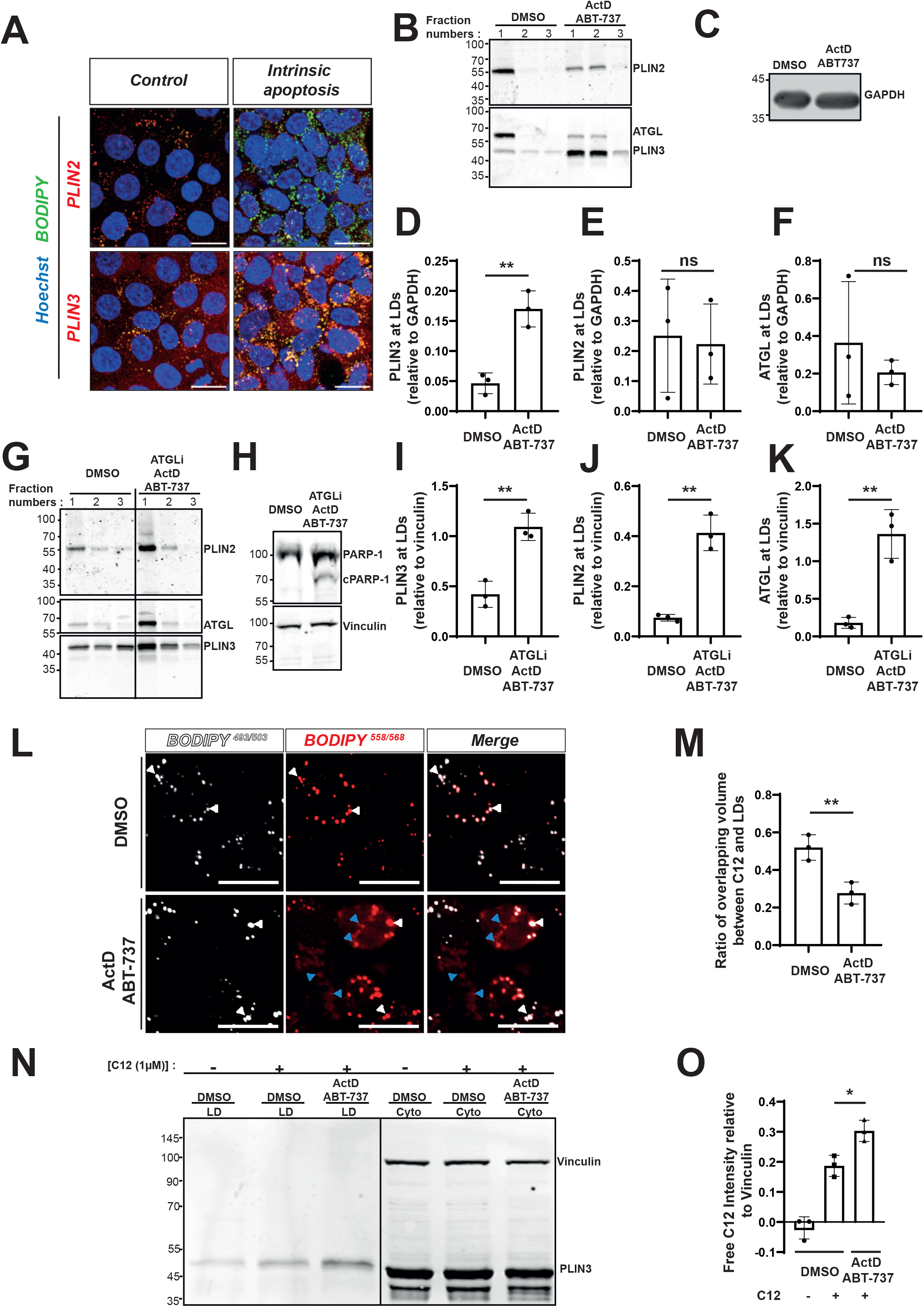
Cell death triggers the redistribution of perilipins and ATGL lipase to promote LD turnover. **(A)** Representative fluorescence images of BxPC3 cells stained for endogenous PLIN2 or PLIN3 (red), LDs (green), and nuclei (blue) 18 h after induction of intrinsic apoptosis by Actinomycin D (ActD, 1 µM) + ABT-737 (1 µM). Scale bars: 20 μm. **(B)** Representative western blot images showing PLIN2, PLIN3, and ATGL levels on the LDs (fractions 1-3) following intrinsic apoptosis induction with Actinomycin D (ActD, 1 µM) + ABT-737 (1 µM) for 5 h. **(C)** LD fractions (1 to 4) were collected, and the rest of the gradient was mixed for GAPDH detection (cell loading control). **(D, E and F)** Quantification of PLIN3, PLIN2 and ATGL at the LDs. Only fractions 1 and 2 were used for densitometric analysis. Data represent the mean ± SD from three independent experiments (N = 3). Statistical analysis was performed using two-tailed unpaired Student’s t-test: **, p < 0.01 **(G)** Representative western blot images showing PLIN2, PLIN3, and ATGL levels on the LDs (fractions 1-3) following inhibition of ATGL activity (with ATGLi, 10 µM) during intrinsic apoptosis (ActD/ABT-737). **(H)** Pooled sucrose gradient fractions excluding the LD fractions (1-4) for Vinculin (cell loading control) and PARP-1 cleavage detection. (**I, J and K**) Quantification of PLIN3, PLIN2 and ATGL at LDs in the presence or absence of ATGLi. Only fractions 1 and 2 were used for densitometric analysis. Data represent the mean ± SD from three independent experiments (N = 3). Statistical analysis was performed using two-tailed unpaired Student’s t-test: **, p < 0.01. **(L)** FA pulse–chase assay. HCT116 WT cells were pulsed overnight with BODIPY^558^/^568^ C12, and fatty-acid localization was analyzed following treatment with ActD/ABT-737 (1 µM each). LDs were labeled with BODIPY^493/503^ (white) and BODIPY^558^/^568^ C12 (red). White arrows indicate regions of colocalization between LDs and BODIPY^558^/^568^ C12. Blue arrows highlight areas where BODIPY^558^/^568^ C12 no longer overlaps with LDs. Scale bars: 20 μm. **(M)** The quantification of BODIPY^558^/^568^ Red C12 volume overlapping with BODIPY^493/503^ was calculated using IMARIS software. Data represent the mean ± SD from three independent experiments (N = 3). Statistical analysis was performed using two-tailed unpaired Student’s t-test: **, p < 0.01. **(N)** Representative western blot analysis of isolated LD fractions and pooled sucrose gradient fractions excluding the LD fractions (Cyto) following overnight pulse labeling with BODIPY^558^/^568^ C12. PLIN3 and Vinculin were used as markers of the LD and cytosolic fractions, respectively. **(O)** Quantification of free BODIPY^558^/^568^ C12 fluorescence in the cytosolic fraction normalized to Vinculin. Data represent the mean ± SD from three independent experiments (N = 3). Statistical analysis was performed using two-tailed unpaired Student’s t-test: *, p < 0.05.

To examine whether the mechanisms underlying LD dynamics during cell death are conserved across species, we next investigated LD turnover *in vivo* during *Drosophila* germ cell death. We examined the localization of perilipins and the *Drosophila* homolog of ATGL, Bmm. Using antibodies against dPlin1/*Lsd-1* and dPlin2/*Lsd-2*—orthologs of human PLIN2, PLIN3, and PLIN5, we found that dPlin1 was not expressed in germ cells (**Fig. S5A**), consistent with its reported expression mainly in fat body tissue ^30, 31^. In contrast, dPlin2-positive LDs were present in healthy germ cells but decreased sharply in dying ones, possibly due to the lipolytic degradation of LDs carrying dPlin2 during germ cell death (**Fig. 3A, B**).

**Fig. 3:**
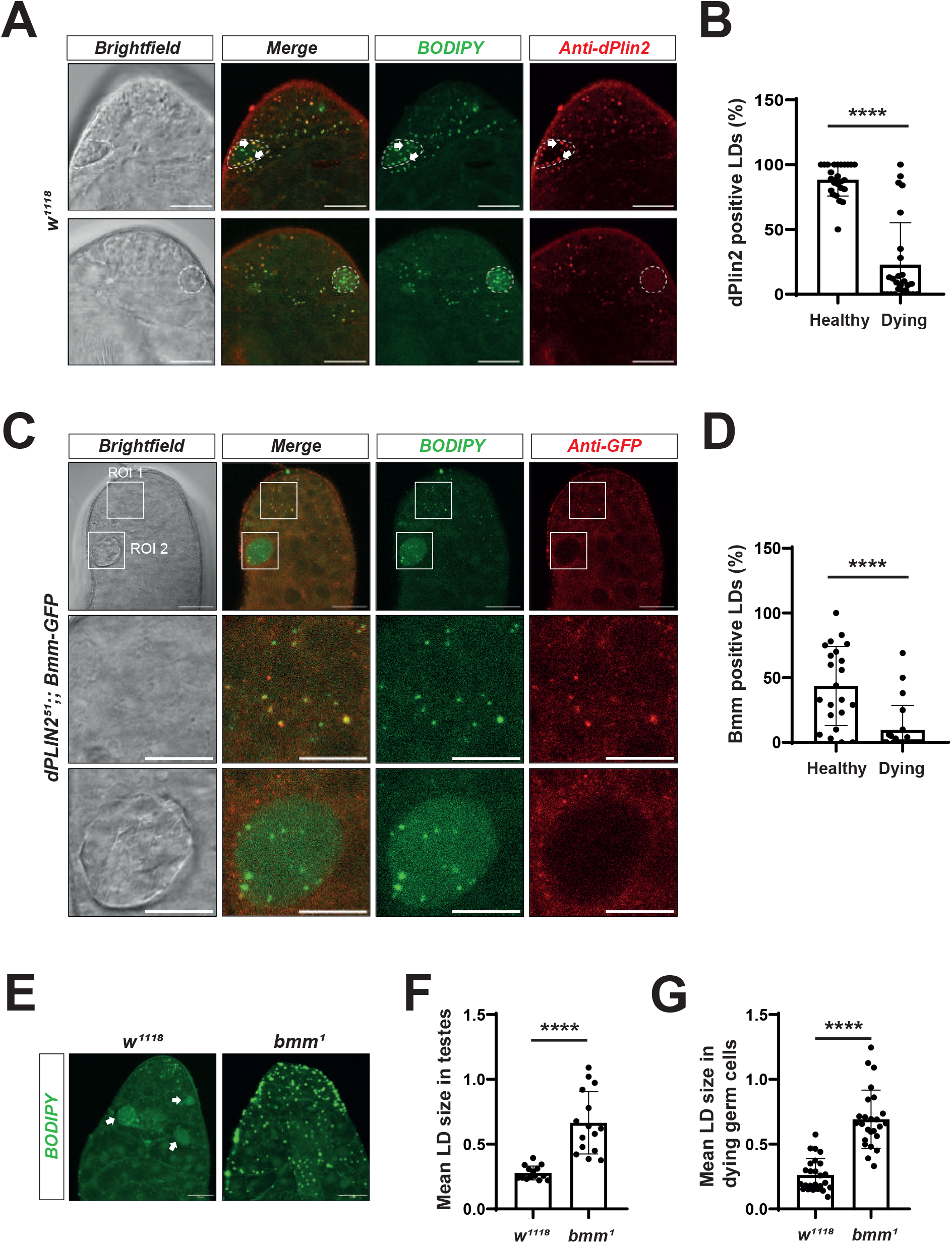
dPlin2 and Bmm redistribution reflects an active lipolysis during germ cell death. **(A)** Representative images of dPlin2 distribution in the apical tip of testes of 5-day-old *w^11^*^18^ flies. Dying germ cells are visualized by brightfield images. The testes were stained with BODPY^493/503^ (green) and antibody against dPlin2 (red). dPlin2 positive LDs (yellow) are indicated with arrows. Dying germ cell are outlined with white dashed lines. Example of dPlin2 positive LDs in dying germ cells is shown in the upper panels. Example of dPlin2 negative LDs in dying germ cells is shown in the lower panels. Scale bars: 20 μm. **(B)** Quantification of the proportion of dPlin2-positive LDs in healthy and dying germ cells. Data represent the mean ± SD from three independent experiments (N = 3, n = 26 for each genotype). Statistical analysis was performed using two-tailed paired Student’s t-test: ****, p < 0.0001. **(C)** Representative images of the apical tip of testes from dPlin2^51^;;Bmm-GFP flies. LDs are labelled with BODIPY^493/503^ (green) and Bmm-GFP using antibody against GFP (red). Regions of interest (ROI) are indicated with solid white squares. ROI 1 indicate regions of healthy germ cells, which are shown at higher magnification in the middle panels. ROI 2 indicate regions of dying germ cells, which are shown at higher magnification in the bottom panels. Scale bars: 20 μm; magnified images: 10 μm. **(D)** Quantification of the proportion of Bmm-GFP positive LDs in healthy and dying germ cells. Data represent the mean ± SD from three independent experiments (N =3, n = 22 for each genotype). Statistical analysis was performed using two-tailed paired Student’s t-test: ****, *p* < 0.0001. **(E)** Representative images of the apical tip of testes from 5-day-old *w^11^*^18^ and *bmm¹* null mutant flies. LDs are stained using BODIPY^493/503^ (green). White arrows indicate dying germ cells. (**F**) Quantification of average LD size per testes in *w^11^*^18^ and *bmm*^1^ flies including both healthy and dying cells. Data represent the mean ± SD of average LD size per testes from three independent experiments (N = 3, each dot represents a single testis; n = 14 and 15 for each genotype, respectively). Statistical analysis was performed using two-tailed unpaired Student’s t-test: ****, *p* < 0.0001. (**G**) Quantification of average LD size in dying germ cells from testes of *w^11^*^18^ and *bmm*^1^ flies. Data represent the mean ± SD of average LD size per testes from three independent experiments (N = 3, each dot represents a single testis; n = 25 for each genotype). Statistical analysis was performed using two-tailed unpaired Student’s t-test: ****, *p* < 0.0001.

We also examined Bmm lipase localization at LDs in germ cells using the Bmm-GFP knock-in line, which showed expected Bmm enrichment around LDs in fat body cells (**Fig. S5B**) ^32^. In contrast, we detected only weak Bmm association with LDs in germ cells (**Fig. S5C**), which could reflect restricted access due to dPlin2 coating. Supporting this, deletion of dPlin2 restored clear Bmm co-localization with LDs in healthy germ cells, (**Fig. 3C**). In addition, *Bmm* positive LDs were reduced in dying germ cells, paralleling the loss of dPlin2, which could be due to increased lipolysis of LDs carrying dPlin2 and Bmm or reduced access of Bmm to LDs (**Fig. 3C, D**).

To directly assess the contribution of Bmm-dependent lipolysis, we compared LD size in dying cells of wild-type testes with those of *bmm*^1^ loss-of-function mutants, in which Bmm-mediated lipolysis is abolished. As expected, *bmm*^1^ mutant testes displayed an overall increase in LD size compared to wild type (**Fig. 3E, F**). Moreover, LDs in dying germ cells of *bmm*^1^ mutants were significantly larger than those in wild-type testes (**Fig. 3G**), indicating that a basal level of Bmm-dependent lipolysis persists in dying germ cells under physiological conditions. In parallel, we investigated whether LD accumulation involves *de novo* lipid synthesis. Dying germ cells display an increased number of LDs compared to healthy cells, suggesting active LD biogenesis (**Fig. 1G**). Consistently, loss-of-function mutants of the DGAT1 ortholog, *midway* (*mdy*) resulted in a reduction in LD size in both healthy and dying germ cells (**Fig. S5D, E**), confirming that TG synthesis is required for LD formation in the testes.

Overall, these findings show that LD accumulation in physiologically dying *Drosophila* germ cells reflects coordinated LD biogenesis alongside sustained lipolytic activity, consistent with ongoing LD turnover. This dual regulation closely mirrors the dynamics observed in apoptotic human cancer cells, supporting a conserved role of LD biogenesis and turnover during regulated cell death across species.

### Fatty acid oxidation has limited contribution on cell death induction

Our results indicate that lipolysis is active both during physiological germ cell death in *Drosophila* and intrinsic apoptosis in human cancer cells. We thus examined the possibility that, a difference in lipolysis might influence the availability of FAs for β-oxidation and energy production, potentially influencing the execution of apoptosis ^33^.

To determine whether mitochondrial β-oxidation contributes to cell death regulation, we first analyzed knock-down of the *Drosophila* ortholog of *carnitine palmitoyltransferase 2 (CPT2)*, the rate-limiting enzyme of β-oxidation in *Drosophila*. Knocking-down *CPT2,* using the early germ cell driver *nos-Gal4,* resulted in loss of germ stem cells preventing measurement of cell death rates in spermatogonia ^34^. To circumvent these developmental defects, *CPT2 RNAi* was instead expressed using the *bam-GAL4* driver, which is expressed at later stage in dividing spermatogonia. Under these conditions, germ stem cell number near the Fas III-positive hub and the frequency of dying spermatogonia were comparable to wild type (**Fig. S6A-C**), indicating that *CPT2* knock-down did not alter physiological germ cell death rates. In BxPC3 and HCT116 cells, we also examined the consequence of inhibiting mitochondrial β-oxidation on intrinsic apoptosis induction using etomoxir, an irreversible CPT1 inhibitor that limits fatty acid entry into mitochondria (**Fig. S6D and E**). We found no significant effect of etomoxir on cell death induction. Collectively these results suggest that fatty-acid oxidation contribution to cell death execution in both human cancer cells and *Drosophila* germ cells is unlikely.

### Protective role of lipid droplets during germ cell death

We next assessed whether altering LD content affects germ cells survival in *Drosophila*. To induce LD accumulation, we used *bmm* mutant testes. A previous report showed that *bmm*^1^ loss of function mutant exhibited an increase in germ stem cell number, a reduction in spermatocyte differentiation and subsequent defects in spermatogenesis, while mitotic divisions of spermatogonial cells appeared unaffected ^24^. We thus assumed that the increased germ stem cell number previously observed would not indirectly affect spermatogonia cell death. We found a reduction of germ cell death in *bmm*^1^ mutant compared to the wild type testes suggesting that the accumulation of LDs inhibits germ cell death **(Fig. 4A, B)**. To further demonstrate the requirement of *bmm* in germ cells, we performed a rescue experiment, in which the re-rexpression of wild-type *bmm* using the *nos* driver restored germ cell death in *bmm*^1^ testes (**Fig. 4C, E**). To confirm that physiological germ cell death is sensitive to LD levels, we tested the effect of LD depletion by overexpressing *bmm.* However, bmm-dependent LD depletion was not very effective possibly due to dPlin2 inhibiting Bmm. To circumvent this, we overexpressed *bmm* in *dplin2* mutant testes, which led to an efficient LD depletion. In this condition we observed an increased germ cell death, further supporting a protective role of LDs in germ cell death (**Fig. 4D, F**). We also used *mdy*-induced LD depletion to evaluate the effect of LDs on cell death. However, we did not observe a difference in germ cell death rates in *mdy* mutant (**Fig. S7A, B**), which was unexpected because *mdy* mutant exhibit increased germ cell apoptosis during oogenesis ^35^. A possible explanation is that *mdy* mutant led to incomplete TG synthesis inhibition as observed with the remaining LDs (**Fig. S5D, E**), which may be due to a role of both Mdy and Dgat2 during spermatogenesis. This, however, could not be assessed because *dgat2* gene is triplicated and no triple KO are available to target its expression efficiently ^26^. Collectively these results show that LD accumulation depending on *bmm* inhibition confers protection to germ cells death.

**Fig. 4.**
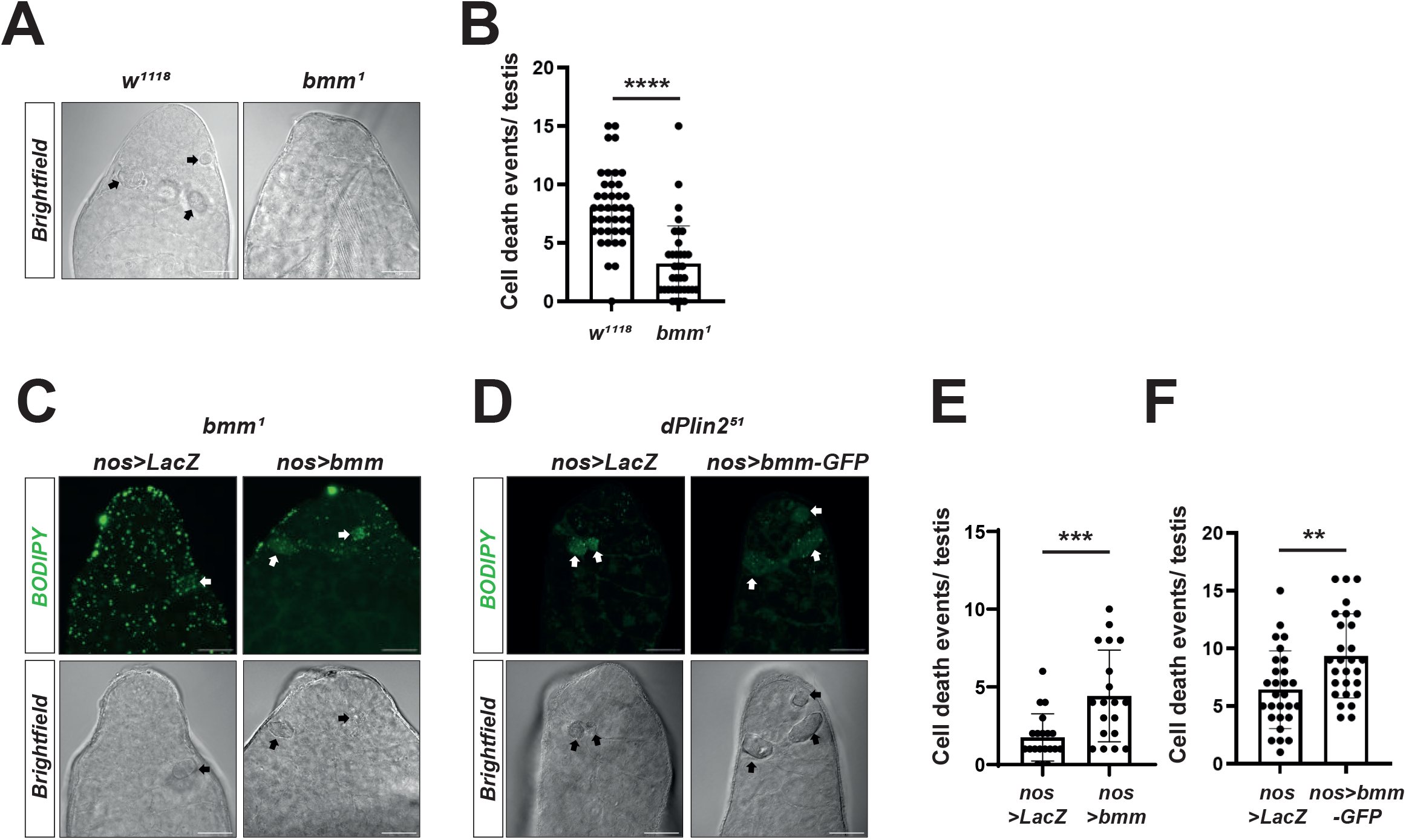
Bmm is cell autonomously required to regulate LD accumulation and germ cell death. **(A)** Representative brightfield images of the apical tip of testes of 5-day-old control (*w^11^*^18^) and *bmm* null mutant flies (*bmm¹)*. Dying germ cells are indicated with black arrows. Scale bars: 20 μm. **(B)** Quantification of the number of dying germ cell in the testes of control (*w^11^*^18^) and *bmm*^1^ flies. Data represent the mean ± SD from three independent experiments (N = 3, each dot represents a single testis; n = 41 and 35 for each genotype, respectively). Statistical analysis was performed using two-tailed unpaired Student’s t-test: ****, *p* < 0.0001. **(C)** Representative images of the apical tip of testes from 5-day-old *bmm*^1^ mutant flies expressing either *LacZ* (control) or *bmm* in early germ cells (*nos-Gal4*). LDs are stained with BODIPY^493/503^ (green). Dying germ cells are indicated with white arrows. Scale bars: 20 μm. **(D)** Representative images of the apical tip of testes from 5-day-old *dPlin2*^51^ mutant flies expressing either *LacZ* (control) or *bmm-GFP* in early germ cells (*nos-Gal4*). LDs were stained with BODIPY^493/503^ (green). Dying germ cells are indicated by white arrows. Scale bars: 20 μm. **(E)** Quantification of the number dying germ cell in the testes depicted in 4C. Data represent the mean ± SD from three independent experiments (N = 3, each dot represents a single testis; n = 20 and 19 for each genotype, respectively). Statistical analysis was performed using two-tailed unpaired Student’s t-test: ***, *p* < 0.001. **(F)** Quantification of the number of dying germ cell in the testes depicted in 4D. Data represent the mean ± SD from three independent experiments (N = 3, each dot represents a single testis; n = 28 for each genotype). Statistical analysis was performed using two-tailed unpaired Student’s t-test: **, *p* < 0.001.

### Dynamic changes in LD protein profile occur during apoptosis

Our results have shown that an active LD turnover is associated with PLIN2, PLIN3 and ATGL redistribution during apoptosis in human cancer cells. We hypothesized that LD protein redistribution may affect the entire LD proteome and possibly cell death regulators, such as the main pro- and anti-apoptotic factors, Bax and Bcl-xL, respectively, which are also reported to bind LDs ^20, 36^. We performed a LD whole proteome analysis using quantitative LC-MS/MS on isolated LDs from apoptotic HCT116 cells. Approximately 2,400 proteins were identified, with 228 showing differential regulation (up/down regulated) (**Fig. 5A and Table S1A, B**). We classified these up- and down-regulated proteins into functional categories that include cellular stress responses, gene expression, lipid metabolism, and DNA repair (**Fig. 5B** and **Table S1C**). These findings indicate that LDs undergo a broad remodeling of their proteome during intrinsic apoptosis, reflecting their active involvement in cellular stress adaptation and metabolic regulation. In agreement with our previous observations, we found an accumulation of PLIN3 on LD fractions in apoptotic HCT116 cells. In addition, we also found Bax and Bcl-xL in the proteome but both remained below the fixed two-fold change threshold across replicates (**Fig. 5A, C)**. To further investigate differences of protein abundance of these cell death regulators, we used a targeted approach by western blotting. While both Bax and Bcl-xL were found at LDs in non-treated cells, we observed a significant increase of Bax and a decrease of Bcl-xL following ActD/ABT-737 treatment (**Fig. 5D-F**), suggesting that a fraction of Bax is relocalized to LDs during the early stage of apoptosis. The increase of Bax in the LD fraction was also observed in HCT116 overexpressing Bax (**Fig. S8A, B**). We attributed the decrease of Bcl-xL on LDs to ABT-737, which binds the BH3 groove of Bcl-xL and disrupts its association with LDs via the V-domain ^37^. Together, these findings reveal that LDs actively engage in the spatial regulation of the pro-apoptotic protein Bax, positioning them as active dynamic modulators of cell death signaling during intrinsic apoptosis.

**Fig. 5.**
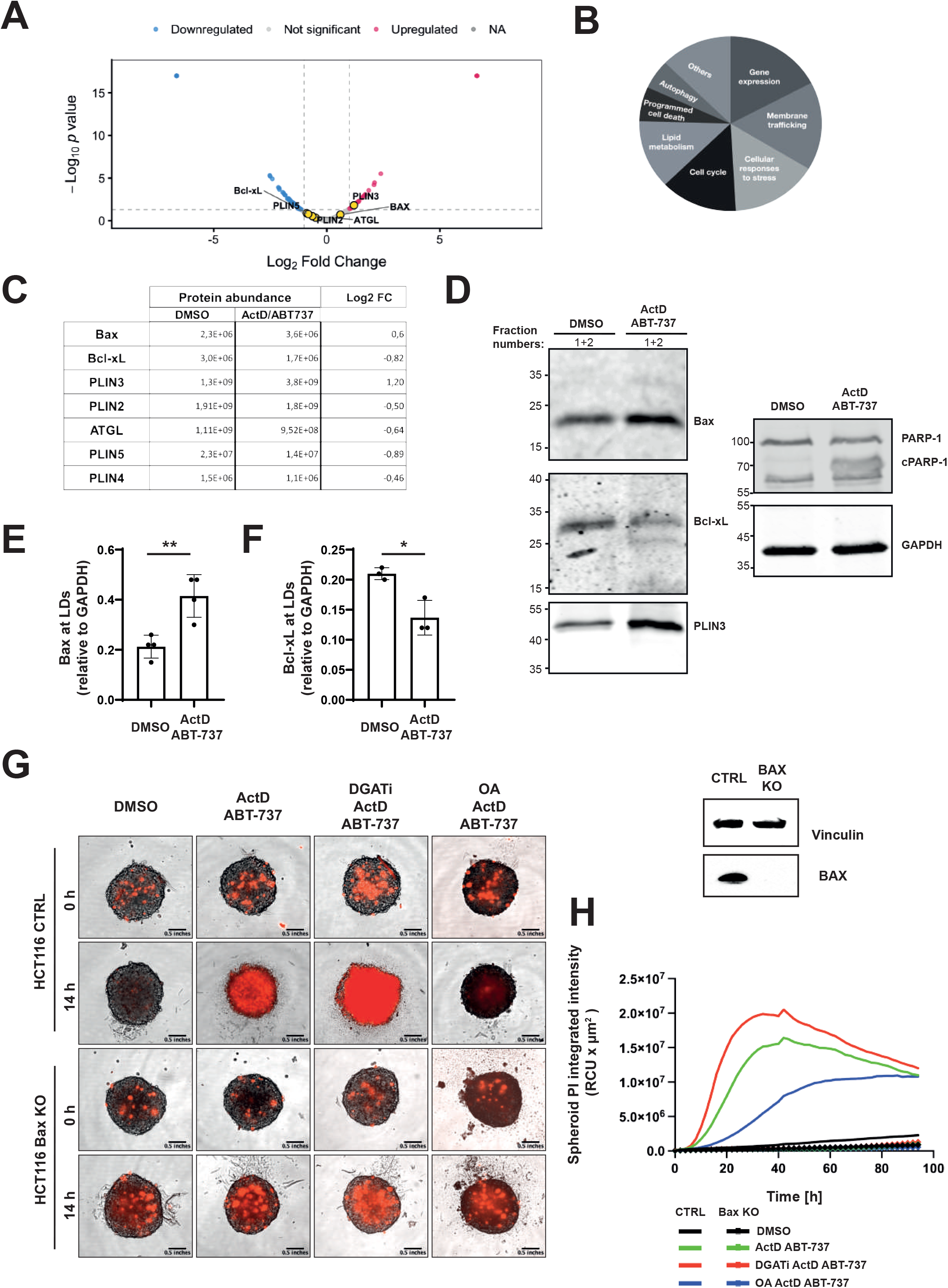
LD proteome remodeling following intrinsic apoptosis-induction in HCT116 cells. **(A)** Volcano plot of proteins detected in isolated LD fractions following intrinsic apoptosis induction in HCT116 cells. Quantitative LC-MS/MS was performed on isolated LD fractions from HCT116 cells treated with ActD/ABT-737 (1 µM each) or DMSO (control) for 5 h. Proteins significantly enriched or depleted in LD fractions upon apoptosis induction are shown in red and blue, respectively. PLIN2, PLIN3, PLIN5, ATGL, Bax, and Bcl-xL are highlighted in yellow. The red and blue dots with value at of −6.64 and +6.64 in log₂ fold change arise from ratios of several proteins whose abundance was zero either before or after ActD/ABT-737 treatment in the initial analysis, respectively. (see Table S1A and B). **(B)** Pie chart showing the distribution of functional classes among proteins with altered abundance in isolated LD fractions following intrinsic apoptosis induction in HCT116 cells (Table S1C). **(C)** Table depicting protein abundance and fold change of the indicated candidate LD proteins. **(D)** Representative western blot images showing Bax, Bcl-xL, PLIN3 levels in the LD fractions following intrinsic apoptosis induction with ActD/ABT-737 (1 µM of each) or DMSO (control) for 5 h. LD fractions (1 to 4) were collected, and the rest of the gradient was mixed for GAPDH (cell loading control) and PARP-1 cleavage detection. **(E, F)** Quantification of Bax and Bcl-xL levels in the LD fraction isolated from cells treated with apoptosis inducers or DMSO control. This experiment is representative of four biological repeats (N = 4). Statistical analysis was performed using two-tailed unpaired Student’s t-test: *, p < 0.05; **, p < 0.01. **(G)** Representative images of HCT116 CTRL and Bax knockout spheroids exposed to the apoptosis inducers ActD/ABT-737, following pretreatment with either oleic acid (100 µM) to promote LD accumulation or a DGAT1/2 inhibitors (15 µM and 10 µM, respectively) to reduce LD levels. On the right: Representative western blot images confirming Bax knockout in HCT116 cells. **(H)** Quantification of spheroid propidium iodide (PI) integrated intensity (RCU × µm²). Data are representative of three independent experiments (N=3) yielding comparable results.

### LDs delay Bax-induced apoptosis

The accumulation of Bax at LDs suggests that LDs may protect against Bax-dependent apoptosis. If so, the extent of Bax-mediated apoptosis should depend on LD abundance. To test this, we induced apoptosis with ActD/ABT-737 under conditions of either LD depletion (via DGAT1/2 inhibition) or LD accumulation (following oleic acid treatment) in wild-type and Bax knockout HCT116 cells. Cells were cultured as 3D spheroids to better recapitulate the tumor architecture and microenvironment and because they exhibit a greater sensitivity to Bax-induced apoptosis than conventional 2D cultures ^38^. We observed that DGAT1/2 inhibition increased apoptosis, whereas oleic acid reduced it as measured by propidium iodide incorporation over 100 h. Consistent with previous findings, ActD/ABT-737-induced apoptosis was Bax-dependent, and completely abolished in Bax KO cells (**Fig. 5G, H**) ^25^. Collectively, these results support a protective role of LDs during apoptosis, likely through partial sequestration of Bax at the LD surface.

### Active Bax translocates from mitochondria to LDs during apoptosis

The inhibition of Bax-dependent apoptosis by LDs suggests that the localization of Bax at LDs correlates with a reduced recruitment at the mitochondria during early apoptosis. To test this, we assessed Bax level on isolated mitochondria from cells in early and late stages of apoptosis. We took advantage that early and late stage of apoptosis could be easily distinguished in HCT116 cells undergoing apoptosis. Indeed, during apoptosis induction, a fraction of cells rounded up and detached while the remaining cells stayed attached. The attached HCT116 cells (AC, early apoptosis) exhibited some but not fully cleaved Caspase 3 processing and were considered as early-stage apoptotic cells. In contrast, the detached cells (floating cells - FC) have high levels of cleaved Caspase 3 and were thus considered as late-stage apoptosis (**Fig. S8C)**. Interestingly, Bax and Bcl-xL, which were found at low levels in untreated or early-stage of apoptosis (AC), increased together with cytochrome-c release at late-stage of apoptosis (FC) in mitochondrial fractions (**Fig. S8D-F)**. These observations support that when Bax is localized at LDs in the early stage of cell death, it is less capable of translocation to mitochondria. Only in the late stages of apoptosis, Bax reaches sufficient levels at the mitochondria.

To investigate the possibility that mitochondrial Bax translocates from mitochondria to LDs through inter-organelle contacts, we examined LD-mitochondria proximity during cell death. Upon ActD/ABT-737 treatment, we observed increased LD/mitochondria proximity by transmission electron microscopy (TEM) and by immunofluorescence using BODIPY^493/503^ and TOM20 to label LDs and mitochondria, respectively **(Fig. 6A-C)**. Using live imaging of HCT116 cells expressing EGFP-Bax, we further detected rapid recruitment of LDs to mitochondria within 30 minutes of treatment with ActD/ABT-737. This was confirmed both by BODIPY^493/503^ staining of LDs and by label-free tomography combined with MitoTracker. During this period, Bax redistributed from a diffuse cytosolic pattern to supramolecular aggregates at mitochondria that were closely apposed to LDs (**Fig. 6D and Fig. S8G**). Together, these observations indicate that LD recruitment to mitochondria accompanies early Bax activation, raising the possibility that Bax activation is required for its association with LDs. This is also supported by the fact that Bax binds LDs via a specific protein domain composed of 2 alpha helices (a5-6), organized in hairpin called V-domain ^20^, which is hidden on the inactive form of the protein ^39^, and exposed only under activation ^40^. To test this, we used active Bax antibody (E4U1V) and observed a fraction of active Bax at LDs after ActD/ABT-737 treatment in HCT116 cells by immuno-TEM and immunofluorescence, respectively (**Fig. 6E, F**). To further investigate whether Bax recruitment to LDs requires Bax activation, we evaluated Bax binding to artificial LDs composed of a triolein neutral lipid core surrounded by a phospholipid monolayer, visualized by Rhodamine-PE fluorescence (**Fig S9A**). Artificial LDs were incubated with cytosolic extracts from HCT116 cells in which Bax activation was induced using the Bax activator BTSA1 ^41^. In this condition we observed a selective accumulation of Bax in the purified LD fraction (**Fig. S9B-E**). Together, these data indicate translocation of active Bax from mitochondria to LDs and imply reduced Bax activation at mitochondria, providing a mechanistic explanation for how LD accumulation delays apoptotic execution (**Fig. 6G**).

**Fig. 6.**
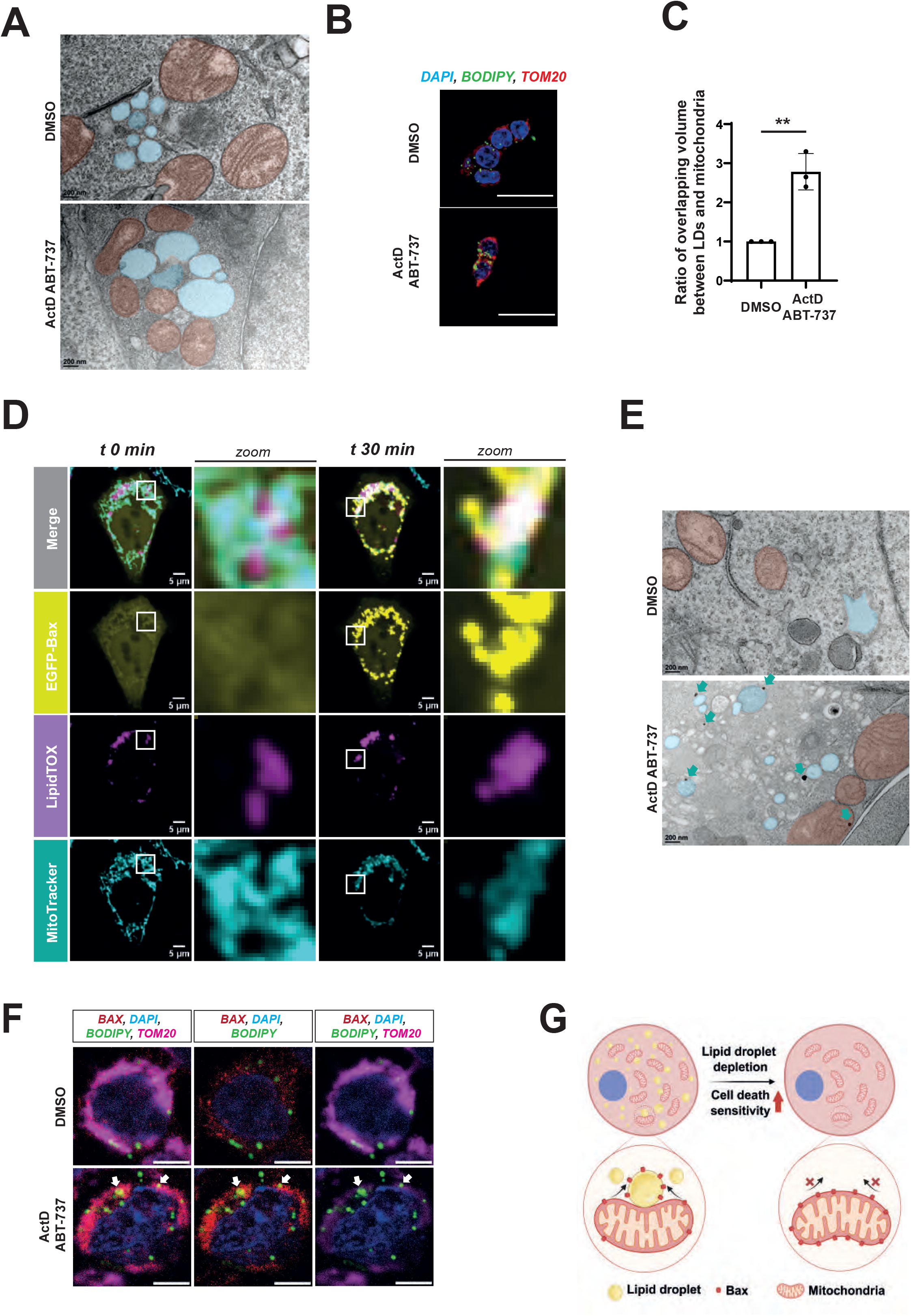
Translocation of activated Bax from mitochondria to LDs delays apoptosis. **(A)** Representative transmission electronic microscopy (TEM) images of HCT116 WT, control (DMSO) or treated with ActD/ABT-737 (1 µM each) for 5 h. Mitochondria and LDs are shown in orange and blue, respectively. Scale bars: 200 nm. **(B)** Representative fluorescence microscopy images of HCT116 WT, control (DMSO) or treated with ActD/ABT-737 (1 µM each) for 5 h. LDs are stained with BODIPY^493/503^ (green), nuclei with DAPI (blue), and mitochondria using an anti-TOM20 antibody (red). Scale bars: 20 μm. **(C)** Quantification of LD and mitochondria contacts, measured as the ratio of overlapping volume between LDs and mitochondria. Data represent the mean ± SD from three independent experiments (N = 3). Statistical analysis was performed using two-tailed unpaired Student’s t-test: **, p < 0.01. **(D)** Representative images of HCT116 cells overexpressing EGFP-Bax treated with ActD/ABT-737 (1 µM each). LipidTOX was used to stain LDs (magenta), MitoTracker to stain mitochondria (cyan) and EGFP-Bax is shown in yellow. Scale bars: 5 μm. **(E)** Representative TEM images showing active Bax localization in HCT116 WT, control (DMSO) or treated with ActD/ABT-737 (1 µM each) for 5 h. Cyan arrows are indicating active Bax revealed by immunogold staining. Mitochondria and LDs are shown in orange and blue, respectively. Scale bars: 200 nm. **(F)** Representative fluorescence microscopy images of HCT116 cells treated with ActD/ABT-737 Cells were stained 5 h after ActD/ABT-737 with BODIPY^493/503^ for LDs (green), anti-TOM20 antibody for mitochondria (blue) and anti-Bax (E4U1V) antibody (which recognize active form of Bax, red). **(G)** Scheme of proposed model in which LDs delay apoptosis by sequestering mitochondrial Bax. During apoptotic stress, LDs accumulate and form increased contacts with mitochondria, enabling the relocalization of Bax from mitochondria to LDs. This redistribution limits Bax accumulation at the mitochondrial outer membrane, thereby restraining mitochondrial outer membrane permeabilization and delaying apoptosis execution. In contrast, LD depletion prevents Bax sequestration, resulting in its retention at mitochondria and increased sensitivity to apoptosis. These findings support a model in which LD accumulation during cell death transiently buffers mitochondrial pro-apoptotic factors, attenuating their activity and delaying apoptosis.

## Discussion

Here, we investigated LD accumulation during cell death and the underlying protective mechanisms in human cancer cells and *Drosophila*. LDs accumulated across diverse apoptotic and non-apoptotic RCD modalities, with necroptosis as a notable exception. This accumulation primarily resulted from *de novo* LD biogenesis, which appears to outpace lipolysis, as lipolytic activity was maintained or even increased during intrinsic apoptosis. In *Drosophila*, loss of the Bmm lipase led to enhanced LD accumulation accompanied by reduced germ cell death, supporting a cytoprotective role for LDs. Dying cells exhibited a marked redistribution of LD-associated proteins, including perilipins and lipases in both *Drosophila* germ cells and human cancer cells undergoing intrinsic apoptosis. In cancer cells, the LD proteome underwent extensive remodeling, incorporating cellular stress response proteins—most notably the pro-apoptotic protein Bax, as confirmed by immunoblotting. Bax enrichment at LDs was facilitated by increased LD–mitochondria contacts, enabling the transfer of its active form. Functionally, sequestration of Bax at LDs delayed the execution of intrinsic apoptosis, establishing LDs as active regulators of cell death signaling through protein redistribution.

### LDs remodel into distinct subpopulations during apoptosis

A previous study showed that intrinsic apoptosis induces *de novo* LD synthesis through upregulating enzymes such as DGAT and Acyl-CoA Synthase (ACS), which redirect fatty acids from mitochondrial β-oxidation toward TG storage ^18^. Our results demonstrate that LD accumulation during intrinsic apoptosis is driven by increased lipid flux, with enhanced *de novo* biogenesis occurring concomitantly with enhanced lipolysis. This reveals that LD metabolism during apoptosis is more dynamic and complex than previously appreciated and gives rise to functionally distinct perilipin-defined LD subpopulations in human cancer cells. PLIN3 specifically accumulates on nascent LDs during early apoptosis, while ATGL loading remains low on these LDs but persists on a PLIN2-positive subpopulation prone to lipolysis, as evidenced by ATGL inhibition experiments and BODIPY C12 pulse-chase assays. This PLIN3-dominated pool expands due to DGAT-dependent TG synthesis, whereas PLIN2-marked LDs undergo active turnover, preventing net lipolysis suppression. Altogether, this dynamic interplay between PLIN3- and PLIN2-containing LDs during apoptosis reflects the emerging view that LD subpopulations play distinct and context-dependent roles in cellular metabolism and fate ^42^.

### LD accumulation during cell death: Conservation Across Species

A LD subpopulation–specific regulation appears broadly conserved between *Drosophila* germ cell death and human cancer cell apoptosis, although with distinct dynamics. In *Drosophila*, the loss of dPlin2- and Bmm-positive LDs during cell death is particularly striking and correlates with sustained lipolysis. Accordingly, *bmm¹* mutants accumulate markedly enlarged LDs in dying germ cells, confirming that Bmm-dependent lipolysis remains active during physiological cell death. In cancer cells, rapid LD turnover obscures similar transitions under normal conditions; however, when *de novo* synthesis is blocked, a comparable pattern emerges, *i.e.* a PLIN2-positive LD population decreasing by lipolysis during apoptosis in HCT116 cells. Thus, both in cancer cells and *Drosophila* germ cells the accumulation of LDs is associated with a marked lipolysis dependent of ATGL/Bmm. Interestingly, while PLIN3-positive LDs are abundantly detected in cancer cells upon apoptosis, accumulating LDs lack dPlin1 or dPlin2 in dying *Drosophila* germ cells. This suggests that an alternative LD protein accumulate at the surface of LDs for their stabilization, as PLIN3 does in human cells. Thus, while the core mechanism of LD remodeling during cell death is conserved across species, the kinetics and visibility of these processes differ according to cellular context.

### The LD Proteome is Dynamically Remodeled During Cell Death

A major advance of our study is the first quantitative LD proteome during apoptosis (∼2,400 proteins identified, 228 differentially regulated), revealing dynamic compositional remodeling as LDs transition from metabolic organelles to protective platforms. PLIN3 enrichment validated *de novo* LD biogenesis, while Bax recruitment highlighted a novel mechanism to delay apoptosis execution. This proteome shift underscores LDs as signaling hubs, adapting to stress.

This dynamic remodeling aligns with emerging evidence across different stress conditions. In neurodegeneration, ApoE4 microglia exhibit distinct LD proteome with enriched vesicle trafficking and depleted β-oxidation proteins, driving excess LD buildup, heightened proinflammatory cytokine release (TNF, IL-1β), and impaired lipid handling that exacerbates AD pathology ^43^. During inflammation, a LD proteome diversification in immune cells occurs, with a marked recruitment of perilipins, signaling kinases, and lipid mediators to orchestrate eicosanoid production, cytokine regulation, and pathogen defense ^44^. In a recent study on rheumatoid arthritis-associated inflammation, tissue-invading CD4^+^ T cells undergo pyroptosis by the pore forming molecule gasdermin D and the acetyltransferase zDHHC5. The authors showed that these two molecules are carried by LDs to the plasma membrane to trigger membrane rupture and cell death ^45^. Our data extend this paradigm, showing that LDs act as carriers of apoptosis regulators and can function as inhibitors of intrinsic apoptosis by partially sequestering active Bax. Collectively, these results highlight that LDs can exert pro- or anti-apoptotic roles, depending on the cellular context.

### LD Sequestration of Activated Bax is a Novel Anti-apoptotic Mechanism

A key discovery from our study is the early accumulation of Bax at LD surface during intrinsic apoptosis, which delays its pore-forming activity at mitochondria. An earlier study has shown that a transfected V-domain of Bax translocates from mitochondria to LDs upon stress in yeast cells or apoptosis in HepG2 cells ^20^. Our data goes beyond these findings by analyzing full length endogenous Bax as well its active form, redistribution at LDs and functional outcome. We showed that Bax redistribution occurs alongside increased LD-mitochondria contacts and translocation of conformationally active Bax, positioning LDs as temporary "traps" for pro-apoptotic executioners. During early apoptosis, Bax localizes primarily to LDs while remaining scarce at mitochondria; only in late-stage detached cells does Bax accumulate at the mitochondria, triggering massive cytochrome c release. This temporal sequestration creates a protective window that functionally delays apoptotic commitment, as evidenced by experiments showing that DGAT inhibition-mediated LD depletion hypersensitizes HCT116 spheroids to Bax-dependent apoptosis whereas oleic acid–induced LD accumulation confers protection, indicating that Bax sequestration at LDs delays apoptotic execution. This positions LDs as adjustable regulators of the apoptosis threshold, capturing cell death effectors to extend cellular survival under stress. Therapeutically, combining DGAT inhibitors with BH3-mimetics may overcome resistance in LD-rich, apoptosis-refractory cancers.

### Mechanisms of LD-Mediated Protection in *Drosophila* Germ Cell Death

Our first evidence for a protective role of LDs comes from the observation in *Drosophila bmm*^1^ mutant, which exhibits both LD accumulation and reduced germ cell death. We further showed that *bmm* is cell autonomously required in germ cells to regulate LD degradation and cell death. Moreover, the overexpression of *bmm*, which depleted LDs, enhanced germ cell death. Collectively these results show that LD accumulation depending on *bmm* inhibition confers protection to regulated germ cell death. An alternative hypothesis is that the lack of lipolysis in *bmm* mutant leads to a subsequent β-oxidation inhibition and reduction of a lipid-dependent energy source required for cell death. Our results showing that CTP2 knock-down does not affect the rate of germ cell death support that fatty-acid oxidation unlikely contribute to cell death in *Drosophila* germ cells. While the precise mechanism underlying LD protection in *Drosophila* germ cell death remains to be elucidated, two hypotheses emerge from our data and the literature. It was proposed that LDs can sequester polyunsaturated fatty acids (PUFAs) from membranes, preventing peroxidation chain reactions during reactive oxygen species (ROS) surges ^12, 17^. In *Drosophila* neural stem cell niches, hypoxia-induced ROS triggers LD biogenesis, protecting cells from PUFA-mediated damage ^14^. Similarly, glial LDs buffer ROS in mitochondrial mutants, averting neurodegeneration ^13, 46^. Critically, dying *Drosophila* germ cells exhibit DHE staining indicative of ROS accumulation ^21^, and *bmm¹* mutants—displaying LD expansion—show reduced death rates. Thus, LDs may protect spermatogonia by neutralizing death-associated ROS, akin to their role in stem cell niches ^14^.

Another plausible mechanism for the protective role of LDs during *Drosophila* germ cell death arises from the observation that LDs capture Bax in human cancer cells, suggesting an analogous sequestration of Bcl-2 family proteins. In *Drosophila*, germ cell death depends on the pro-apoptotic Bcl-2 homologs Debcl and Buffy ^21, 22^, raising the possibility that LDs similarly sequester Debcl/Buffy and thereby delay mitochondrial commitment to apoptosis. Testing this hypothesis will require dedicated localization studies, which in turn necessitate the generation of appropriate genetic and imaging tools. These non-mutually exclusive mechanisms—antioxidant buffering and effector sequestration—likely act together to protect dying germ cells, mirroring LD-mediated protection in cancer cells while being tuned to physiological contexts.

## Material and Methods

### Fly Stocks

Flies were maintained on standard yeast medium at 25°C unless otherwise noted. The following stocks were obtained from the Bloomington Drosophila Stock Center: *w^11^*^18^ were used as control flies, *nos-GAL4* (BL7415), *bam-GAL4* (BL80579), *C587-GAL4* (BL67747), *UAS-LacZ* (BL1776)*, UAS-LacZ* (BL1777), *UAS-RFP-NLS* (BL30556), *UAS-mCD8::GFP* (BL32186), *UAS-Lam::GFP* (BL7376), *UAS-bmm::EGFP* (BL98110), *vas-AID::EGFP* (BL76126), *bmm::GFP* (BL94600), *mdy^QX^*^25^ (BL5095), *Lsd-2*^51^ (BL24336), *UAS-GFP RNAi* (BL9331), *UAS-cpt2 RNAi* (BL51900). The *UAS-LacZ RNAi* (VDRC 51446) was obtained from the Vienna Drosophila Resource Center. The *bmm*^1^ and *UAS-bmm* were kindly provided by Ronald P Kühnlein (University of Graz). The *UAS-actDrice* fly line is described in ^47^.

### LysoTracker staining of *Drosophila* testes

Whole-mount testes were dissected from adult *Drosophila* in Schneider’s medium (Sigma-Aldrich, S0146) and incubated with LysoTracker Red DND-99 (Thermo Fisher Scientific, L7528, 1:500 dilution in Schneider’s medium) for 30 min at room temperature (RT). Samples were then rinsed twice with Schneider’s medium for 10 min each. Finally, testes were either mounted in Vectashield mounting medium containing DAPI (Vector Laboratories, H-2000-10) or fixed for 30 min in 4% paraformaldehyde (Electronic Microscopy Science, #15710) for subsequent staining.

### Whole mount propidium iodide staining of *Drosophila* testes

Whole-mount testes were dissected from adult *Drosophila* in Schneider’s medium and incubated with propidium iodide (Sigma-Aldrich, P4864, 50 μg/ml in Schneider’s medium) for 30 min at RT. Samples were then rinsed three times with Schneider’s medium for 10 min each. Finally, testes were fixed for 30 min in 4% PFA for subsequent staining.

### Immunofluorescence staining of *Drosophila* testes

Testes were dissected in Schneider’s medium and fixed in 4% PFA for 30 min. After fixation, samples were rinsed three times for 10 min each with PBST (0.1% Triton X-100 in PBS). Samples were blocked using 5% bovine serum albumin (BSA, Euromedex, #04-100-812-E) in PBST for 40 min at room temperature and incubated with the appropriate primary antibody overnight at 4°C. Samples were then rinsed three times for 10 min each with PBST at room temperature. Following the incubation with appropriate Alexa Fluor-conjugated secondary antibodies (Invitrogen, 1:400) for 2 hours at RT. Finally, samples were rinsed 3 times again in PBST and mounted in Vectashield with DAPI (Vector Laboratories, H-2000-10).

Primary antibodies used included: mouse anti-GFP (Thermo Fisher Scientific, A11120, 1:400); mouse anti-Fasciclin III (DSHB, 7G10, 1:400); rabbit anti-dPlin1 (1:3000) and rabbit anti-dPlin2 (1:3000), were gifts from Ronald P Kühnlein (University of Graz); anti-Traffic jam (1:3000) was a gift from Eli Arama (Weizmann Institute of Science). Secondary antibodies were Alexa Fluor-conjugated and used at 1:400 dilution. Secondary antibodies used in this study were goat anti-guinea pig Alexa 633 (1:400; Invitrogen, A-21105), goat anti-rabbit Alexa 568 (1:400; Invitrogen, A-11011), and goat anti-mouse Alexa 546 (1:400; Invitrogen, A-21133).

### LD staining in *Drosophila* testes

Testes were rinsed three times for 10 min each with PBST after fixation or secondary antibody staining. Then the samples were incubated with either BODIPY^493/503^ (Thermo Fisher Scientific, D3922, 5 µg/mL in PBST) overnight at 4°C or LipidTOX (Thermo Fisher Scientific, H34477, 1:500 in PBS) for 40 min at room temperature. After BODIPY staining, samples were rinsed three times for 10 min each, while after LipidTOX staining, samples were rinsed only once (2 min each). All sections were mounted in Vectashield with DAPI. All images of testes were acquired as multiple sections (*z*-stacks) using a Zeiss LSM 800/980 Laser Scanning confocal microscope. The image processing and LD measurement were executed using ImageJ ^48^.

### LD quantification in fly testes

The LD quantification was performed as previously described ^13^. Images were first filtered for noise using Gaussian Blur 3D (σ = 1), after which Z-projections of every 6 slices (5 µm) in one Z-stack were generated for each testis. The MaxEntropy thresholding algorithm was then used for identification of LD number and area. The program was applied for all the Z-projections in each testis.

### Cell culture

BxPC3 cells were cultured in RPMI 1640 medium (Thermo Fisher Scientific, 61870-044), supplemented with 10% fetal bovine serum (FBS; Eurobio, CVFSVF00-01) and antibiotics (penicillin/streptomycin; Thermo Fisher Scientific, 15140-122).

HCT116 cells were cultured in high-glucose Dulbecco’s Modified Eagle Medium (DMEM; Thermo Fisher Scientific, 61965-059), supplemented with 10% FBS (Eurobio, CVFSVF00-01), 2 mM L-glutamine (Thermo Fisher Scientific, 25030-024), 1 mM sodium pyruvate (Thermo Fisher Scientific, 11360-039), non-essential amino acids (Thermo Fisher Scientific, 11140-035), and antibiotics (penicillin/streptomycin; Thermo Fisher Scientific, 15140-122).

Cells were passaged every 3–4 days and regularly checked for mycoplasma contamination using the MycoAlert™ Mycoplasma Detection Kit (Lonza, LT07-318). Where applicable, cells were treated with following compounds:

ATGLi/NG497 (MedChemExpress, HY-148756), actinomycin D (Sigma-Aldrich, 9415), ABT-737 (Clinisciences, A8193), cyclohexamide (Sigma-Aldrich, C7698), human TNF alpha (Roche, 11371843001), Z-VAD-FMK (Clinisciences, A12373), BV6 (Invivogen, inh-bv6), doxycycline (Sigma-Aldrich, D9891), ionomycin (Sigma-Aldrich, I0634), DGAT1i/A922500 (MedChemExpress, HY-10038), DGAT2i/PF-06424439 (MedChemExpress, HY-108341A), Etomoxir (MedChemExpress, HY-50202A), Q-VD-OPh (Clinisciences, HY-12305), oleic acid (Sigma-Aldrich, O3008), BTSA1 (MedChem HY-123054).

### Generation of Bax overexpressing cells

Parental cells were co-transfected with a plasmid encoding the SB transposase^40^ and either an empty vector (EV) or plasmid with Bax coding sequence, using Lipofectamine 2000 (Thermo Fisher Scientific, 11668019).

At 72 h post-transfection, cells were subjected to antibiotic selection: 500 µg/mL Zeocin (InvivoGen, ant-zn) for Bax overexpressing (OE) cells. Stably transfected populations were subsequently sorted using a FACSAria cell sorter (BD Biosciences) based on constitutive mCherry (Bax OE), to obtain homogeneous populations. For overexpression of transgenes doxycycline (Sigma-Aldrich, D9891) in concentration 0.1-1.0 µg/mL was used.

### CRISPR/Cas9-mediated Bax knockout

Bax sgRNA was inserted into the LentiCRISPRv2-Blast vector (Addgene plasmid #83480) according to the protocol described by ^49^. Stable knockout cell populations were generated by lentiviral transduction followed by selection with 10 μg/mL blasticidin (InvivoGen, ant-bl). The following sgRNA oligonucleotides were used for CRISPR/Cas9-mediated gene knockout: Forward, 5′-CACCGAGTAGAAAAGGGCGACAACC-3′; Reverse, 5′-AAACGGTTGTCGCCCTTTTCTACTC-3′.

### LD quantification in cancer cells

Cancer cells were seeded into 96-well black PhenoPlates (PerkinElmer, 6055302). To visualize and quantify LD number and size, cells were stained with BODIPY^493/503^ (0.5 µg/mL; Thermo Fisher Scientific, D3922) or LipidTOX (1:1000; Thermo Fisher Scientific, H34475), together with Hoechst 33342 (1 µg/mL; Biotium, 3656055) for nuclear labeling and cell counting. Staining was performed at 37°C for 30 min, followed by imaging with the Opera Phenix Plus high-content confocal system (Revvity). Counting was performed by Harmony™ software (Revvity).

### Immunofluorescence

Cancer cells (1 × 10^4^ or 1.5 × 10^4^ cells per well) were seeded in black 96-well PhenoPlates (PerkinElmer, 6055302). Cells were fixed with 4% paraformaldehyde (PFA; Euromedex, 15714) for 15 min at RT, then washed three times with PBS. Permeabilization and blocking were performed using 3% BSA and 0.1% Triton X-100 in PBS for 45 min.

Cells were incubated for 1 h at room temperature with primary antibodies (1:300): ADRP/Perilipin 2 (Proteintech, 15294-1-AP), ATGL (Cell Signaling, 2138), Bax (Cell Signaling, 41162), Bcl-xL (Cell Signaling, 2764), and TIP47/Perilipin 3 (Proteintech, 15694-1-AP). After three washes in 0.1% Triton X-100 in PBS, secondary Alexa Fluor-conjugated antibodies (1:1000; Thermo Fisher Scientific, A11011, A11004, A21235, A21244, A31553) were applied together with BODIPY^493/503^ (0.5 µg/mL; Thermo Fisher Scientific, D3922) and Hoechst 33342 (1 µg/mL; Biotium, 3656055) for 1 h. Cells were washed three times with PBS and imaged using the Opera Phenix Plus high-content confocal system (Revvity). Counting was performed by Harmony™ software (Revvity). For active Bax detection, a modified protocol was used. Permeabilization was performed with CHAPS-based buffer instead of Triton X-100, and PBS was used for all washing steps.

### In real-time imaging of EGFP-BAX during cell death

HCT116 cells were seeded into black 96-well PhenoPlates (PerkinElmer, 6055302) at a density of 1.5 × 10^4^ cells per well and transfected with EGFP- Bax plasmid using Lipofectamine 2000 (Thermo Fisher Scientific, 11668019) according to the manufacturer’s instructions. Approximately 7 h after transfection, cells were stained with HCS LipidTOX Deep Red (1:1000; Thermo Fisher Scientific, H34477) and MitoTracker™ Red CMXRos (1:10,000; Thermo Fisher Scientific, M7512) for 30 min at 37 °C. Following staining, live-cell time-lapse imaging was performed using an Opera Phenix Plus high-content confocal imaging system (Revvity), with images acquired every 5 min.

For holotomographic imaging, EGFP- Bax-transfected cells were stained only with MitoTracker™ Red CMXRos (1:10,000) and imaged using a 3D Cell-Explorer Fluo system (Nanolive), with images acquired every 5 min.

### Spheroid formation

HCT-116 spheroids were generated by preparing a cell suspension at a density of 5×10^5^ cells/mL and seeding 100 µL into each well of an ultra-low attachment, round-bottom 96-well plate (Corning, 7007), resulting in 5×10^4^ cells per well. The plates were incubated under standard culture conditions for 3 days to allow the formation of compact spheroids. Once mature, the spheroids were used for subsequent experiments.

### Cell death assay using IncuCyte Imager

Cancer cells (1 × 10^4^ or 1.5 × 10^4^ cells per well) were seeded in 96-well plates. After 24 h, cells were treated with the indicated compounds to induce cell death. To label dead cells, the culture medium was supplemented with Sytox Green (30 nM; Life Technologies, 1846592). Alternatively, spheroids formed in ultra-low attachment, round-bottom 96-well plates (Corning, 7007) were stained with propidium iodide (3.3 µg/mL; Sigma-Aldrich, P4864). Cells cultured in two-dimensional (2D) or three-dimensional (3D) conditions were imaged at 60 min intervals using the IncuCyte live-cell imaging system (Sartorius).

### LD isolation

LDs isolation was performed as previously described ^48^. Briefly, cells were grown on 15-cm dishes till approximately 80% of confluence then treated with indicated drugs as described in figure legends. Cells were rinsed with 10 mL of TNE (20 mM Tris/HCl, pH 8, 130 mM NaCl). Then, cells were lysed in 1 mL of TNE supplemented with protease inhibitor Complete (Roche, 04693132001) using a syringe (U-100 insulin 1mL 0.33 mm × 12.7 mm). Cell debris were pelleted at 1,500 × g for 3 min at 4°C. The cell lysate was transferred to a 13 mL ultracentrifuge tube (Beckman, 9/16 × 3 1/2 UC tube) and the sucrose concentration in TNE was adjusted to 0.43 M. The samples were then overlaid with 2 mL of 0.27 mM Sucrose, 2 mL of 0.135 mM sucrose and 2 mL of TNE buffer. LD isolation was performed by ultracentrifugation at 29,600 rpm for 1 h at 4°C (Beckman, SW60Ti). LD fractions were manually collected and analyzed.

### Artificial LD formation

Artificial lipid droplets (ALDs) were prepared as previously described ^50^. Briefly, 1 mg triolein (Sigma-Aldrich, T7140-500MG) was mixed with 10 mg total phospholipids (Avanti Polar Lipids) composed of 60% POPC, 25% POPE, 10% POPS, and 5% Rhodamine-PE (v/v). Lipids dissolved in chloroform were combined in a glass tube, and the solvent was evaporated under a gentle stream of nitrogen until a dry lipid film was obtained (∼1 h). The lipid film was hydrated with 1 mL LD buffer (20 mM HEPES, 100 mM KCl, 2 mM MgCl₂, pH 7.4) and vortexed vigorously for 1 min to generate a crude emulsion.

The emulsion was subsequently sonicated for seven cycles of 10 s at 40% amplitude, with cooling between cycles to prevent overheating, to reduce the size of the artificial LDs. The preparation was then centrifuged at 1,500 × g for 5 min to pellet insoluble lipid aggregates. The floating artificial lipid droplets were carefully collected and used immediately for subsequent experiments. For confocal imaging, glass coverslips were prepared as previously described ^51^.

### BAX recruitment assay

To assess Bax recruitment, artificial lipid droplets (ALDs) were stained with BODIPY^493/503^ and divided into two fractions. The ALD fractions were then incubated with cytosolic extracts obtained from HCT116 cells following the isolation of endogenous lipid droplets. Bax activation was induced by the addition of 1 µM BTSA1, and the mixtures were incubated at room temperature for 2 h. Following incubation, ALDs were re-isolated by flotation using the protocol described above. The ALD fractions were collected, and the BODIPY^493/503^ fluorescence intensity was measured to quantify LD recovery. Recruitment of proteins to aLDs was subsequently analyzed by western blot.

### Mass spectrometry

Sample preparation: Samples were processed using SP3 beads following the protocol described ^52^. Briefly, samples were reduced (0.25 M TCEP, 58°C, 45 min) and alkylated (0.5 M iodoacetamide, RT, 45 min). SP3 magnetic beads at 50 µg/µL were then added to the samples using a bead-to-protein ratio of 10. After ethanol addition, samples were incubated for 5 min at 1,000 rpm at room temperature to allow protein binding.

Following three washes of the beads with 80% ethanol, the beads were resuspended in 96 µL of 100 mM ammonium bicarbonate, and 2–4 µL of a LysC/Trypsin enzyme mixture (0.1 µg/µL) were added. Digestion was carried out for 18 h at 37°C with shaking at 1000 rpm.

After bead centrifugation (20,000 g, 2 min), supernatants were collected, dried, and resuspended in 0.1% formic acid.

LC–MS/MS analysis: After peptide quantification (Pierce Fluorometric Peptide Assay, Thermo Fisher Scientific), an amount corresponding to 300 ng of peptides per sample was injected into an Exploris 480 mass spectrometer operating in DIA HCD mode using a 60 min chromatographic gradient.

Raw data files were processed using Proteome Discoverer 3.2 with the CHIMERYS search engine, using the Human SwissProt database together with a contaminant database (commonly observed contaminant proteins in proteomics analyses). Peptide and protein identifications were validated using a false discovery rate (FDR) threshold of 1%.

Quantification was performed in DIA mode using MS2 fragment ions derived from the MS/MS analysis of peptide precursor ions. The quantification ratio of the LD proteome from HCT116 cells treated with ActD and ABT-737 versus DMSO-treated controls was calculated after averaging biological replicates. Protein ratios were statistically evaluated using a pairwise ratio approach followed by a t-test. By convention, proteins were considered upregulated or downregulated when the quantification ratio was >2 or <0.5, respectively, with an associated p-value <0.05.

### Crude mitochondria isolation

Crude mitochondria isolation and cytochrome c release assays were performed as previously described ^53^. Cells were grown till approximately 80% of confluence. Then treated with indicated drugs as described in figure legends. Cells were washed with 10 mL PBS and mechanically detached from the plate. Cells were then centrifuged (800 × g at 4°C for 5 min). The cell pellet was resuspended in 500 μL MB buffer (210 mM mannitol, 70 mM saccharose, 1 mM EDTA and 10 mM Hepes pH 7.7) supplemented with protease inhibitor cocktail (Sigma-Aldrich, 4693116001). Cells were homogenized using insulin syringes (U-100 INSULIN 1mL 0.33 mm ×x 12.7 mm). Cell debris were removed by centrifugation (600 × g at 4°C for 5 min). The supernatant was recentrifuged (800 × g at 4°C for 5 min). Mitochondria were isolated by centrifugation at 8,000 × g for 10 min at 4°C. The mitochondrial pellet was then resuspended in 200 μL of MB buffer, made up to 2 mL and centrifuged again (8,000 × g for 10 min at 4°C). The mitochondrial pellet was resuspended in 200 μL of MB buffer, used directly for Western Blot analysis.

### Protein Extraction and Western Blot Analysis

Total protein extraction was performed using RIPA buffer supplemented with a protease inhibitor cocktail (Sigma-Aldrich, 4693116001) and phosphatase inhibitors (Sigma-Aldrich, 04906845001). Total protein concentration was measured by Bradford method using the Protein Assay Dye Reagent Concentrate (Bio-Rad, 5000006). Equal amounts of protein (10-20 µg per sample) were separated by SDS-PAGE in TG-SDS buffer (Euromedex, EU0510-B) and transferred onto nitrocellulose membranes using the TransBlot Turbo Transfer System (Bio-Rad, 1704150EDU).

Membranes were blocked for 1 h at room temperature with 5% BSA dissolved in TBS containing 0.1% Tween-20 (TBS-T) to reduce non-specific binding. After blocking, membranes were incubated overnight at 4°C with primary antibodies diluted 1:1000 in 5% BSA in TBS-T.

The following primary antibodies were used: ADRP/Perilipin 2 (Proteintech, 15294-1-AP), ATGL (Cell Signaling, 2138), Bax (Cell Signaling, 41162), Bcl-xL (Cell Signaling, 2764), Cleaved caspase 3 (Cell Signaling, 9664), Cleaved PARP (Cell Signaling, 9532), COX IV (Cell Signaling, 4950), Cytochrome c (Cell Signaling, 12963, 11940), TOM20 (sc-17764 Santa Cruz), GAPDH (Cell Signaling, 97166), Pro-caspase 3 (Cell Signaling, 9662), TIP47/Perilipin 3 (Proteintech, 15694-1-AP), Vinculin (Sigma-Aldrich, V9131).

After washing the membranes three times with TBS-T, membranes were incubated with species-appropriate secondary antibodies conjugated to infrared fluorescent dyes (Li-Cor Biosciences, 926-68070 or 926-32211). Following another series of washes, signals were visualized using the Odyssey Clx Infrared Imaging System (Li-Cor Biosciences). Densitometric analysis was performed using Image Studio Lite software (Li-Cor Biosciences).

### Real-Time Quantitative Reverse Transcription PCR (RT-qPCR)

Total RNA was extracted via NucleoSpin RNA Kit (Macherey Nagel, 740955) according to the manufacturer’s protocol. For cDNA synthesis, 0.5 µg of total RNA was reverse transcribed using the Sensifast cDNA synthesis kit (Bioline, BIO-65053).

Primers targeting specific genes were generated using NCBI’s Primer-BLAST design tool and are detailed in Table 1. *HPRT* served as the internal control for normalization. The thermal cycling protocol for qRT-PCR involved an initial enzyme activation at 95°C for 2 min, followed by 40 amplification cycles consisting of denaturation at 95°C for 5 s and combined annealing/extension steps at 60°C and 72°C for 30 s each.

**Table 1.**
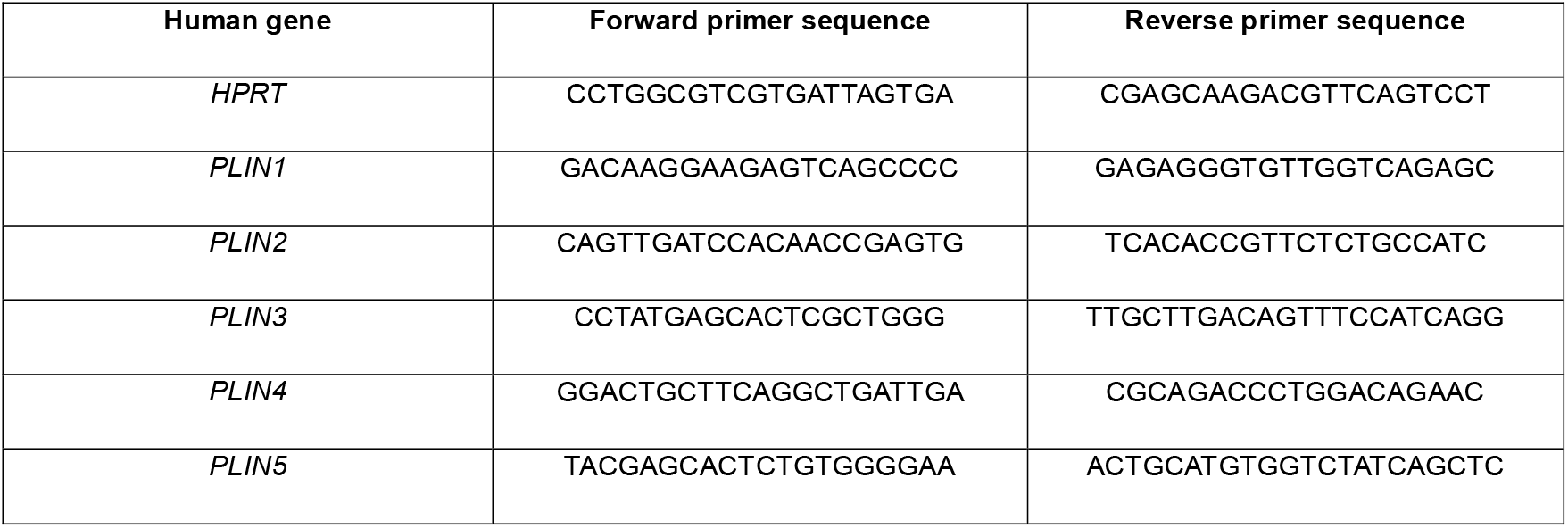

### Transmission electron microscopy

For ultrastructural study, 1 ×10^6^ HCT116 WT cells were treated with 1 µM of ActD and ABT-737 and fixed after 5 h with 2% glutaraldehyde (EMS) in 0.1 M sodium cacodylate (pH 7.4) buffer at 4°C. After washing three times in 0.2 M sodium cacodylate buffer, cell cultures were post-fixed with 2% osmium tetroxide (EMS) at room temperature for 1 h and dehydrated in a graded series of ethanol at room temperature and embedded in Epon. After polymerization, ultrathin sections (100 nm) were cut on a UC7(Leica) ultramicrotome and collected on 200 mesh grids. Sections were stained with uranyl acetate and lead citrate before observations on a Jeol 1400JEM (Tokyo, Japan) transmission electron microscope equipped with a camera in bottom position Orius 1000 (Gaëtan Ametek) and Digital Micrograph (Product version 1.7). This microscope is located at CIQLE (Centre d’Imagerie Quantitative Lyon Est), platform UCBL.

For Active Bax immunogold TEM, HCT116 cells were fixed with 2% glutaraldehyde (EMS) in 0.1 M sodium cacodylate (pH 7.4) buffer at room temperature. After washing three times in 0.2 M sodium cacodylate buffer, HCT116 cells were post-fixed with 1% aqueous osmium tetroxide (EMS) at room temperature for 1 h and contrasted with tea Oolong in 0.2 M pH 7.4 cacodylate (EM grade) for 1 h at room temperature. After washing three times in 0.2 M sodium cacodylate and 1 time in water, cells were dehydrated in a graded series of ethanol at room temperature and embedded in Epon. After polymerization, ultrathin sections (100 nm) were cut on a UC7 (Leica) ultramicrotome and collected on nickel grids (150 mesh).

Immunogold labelling was performed by floating the grids on drops of reactive media. A first step of unmaking the epitope is carried out with metaperiodate 12.5%. Sections were successively washed four times in water. Then, nonspecific sites were coated with 1% BSA and 1% normal goat serum in 50 mM Tris-HCl, pH 7.4, for 2 h at RT. Thereafter, incubation was carried out overnight at 4°C in a wet chamber with the primary antibody. Sections were successively washed three times in 50 mM Tris-HCl, pH 7.4 and pH 8.2 at RT. Nonspecific sites were coated with 1% BSA and 1% normal goat serum in 50 mM Tris-HCl, pH 8.2, for 1.5 h at RT. They were transferred in a wet chamber for 1.5 h at RT in 1% BSA, 50 mM Tris-HCl, pH 8.2 for 1.5 h at RT, labelled with gold-conjugated secondary antibody (Aurion). Sections were successively washed three times in 50 mM Tris-HCl pH pH 8.2 and pH 7.4 and three times infiltrated with distilled water. The immunocomplex was fixed by a wash in 4% glutaraldehyde for 3 min. Sections were stained with lead citrate for 5 min and observed with a transmission electron microscope JEOL 1400JEM (Tokyo, Japan) operating at 100 kV equipped with a camera Orius 1000 gatan and Digital Micrograph, product version: 1.7. This microscope is located at CIQLE (Centre d’Imagerie Quantitative Lyon Est), platform UCBL.

### Fatty acid pulse-chase assay

The fatty acid pulse-chase assay was performed as previously described ^54^. 15 × 10^3^ HCT116 WT were plated on a 96-well PhenoPlate (Revvity, 8793-25341). Cells were treated with 1 µM BODIPY Red C12 (Invitrogen, BODIPY^558^/^568^ C12 2896569) overnight. Cells were then washed three times with complete media and chased for 5 h in the same media complemented with 1 µM of each ActD/ABT-737. Cells were fixed after 5 h of treatment with 4% PFA (PFA; Euromedex, 15714), and the immunostaining was performed as previously described. Total LD were labelled using green BODIPY^493/503^ (0.5 µg/mL; Thermo Fisher Scientific, D3922), and mitochondria were labelled using TOM20 (sc-17764 Santa Cruz) antibody.

### Statistical analyses

Statistical analyses were performed using GraphPad Prism (v8.0.1). Data are presented as individual values with mean ± SD. Normality and homogeneity of variance were assessed using the Shapiro–Wilk and Brown–Forsythe tests, respectively. Based on these results, appropriate parametric or nonparametric tests (t-test or ANOVA) were applied. A p-value < 0.05 was considered statistically significant (*p < 0.05, **p < 0.01, ***p < 0.001). Details of statistical tests are provided in the figure legends.

## Acknowledgments

We thank our colleagues from the fly community and fly stock centers for providing fly stocks and antibodies used in this study. We thank the imaging LYMIC-PLATIM, the fly ARTHRO-TOOLS and Protein Science facilities (SFR Biosciences, Lyon) as well as CIQLE (Centre d’Imagerie Quantitative Lyon Est), platform UCBL. Biorender was used to generate figure model.

This work was supported by INCA PLBIO23-188 to BM, GI and NA, Ligue contre le Cancer Loire (CD42) and Saone et Loire (CD71) to BM, GI and KS, French National Research Agency as part of the “Investissements d’Avenir ExcellencES” program from France 2030 (SHAPE-Med@Lyon; ANR-22-EXES-0012) for GI and KS, and LabEX DEVweCAN for GI and KS. YS benefited from a PhD fellowship from the the program of China Scholarship Council (CSC).

## Declaration of generative AI and AI-assisted technologies in the manuscript preparation process

During the preparation of this work the author(s) used Perplexity AI in order to improve English writing in some sections of the manuscript. After using this tool/service, the author(s) reviewed and edited the content as needed and take(s) full responsibility for the content of the published article.

## References

1. Mathiowetz, A.J. & Olzmann, J.A. Lipid droplets and cellular lipid flux. Nat Cell Biol 26, 331–345 (2024).

2. Fanning, S., Selkoe, D. & Dettmer, U. Parkinson’s disease: proteinopathy or lipidopathy? NPJ Parkinson’s disease 6, 3 (2020).

3. Krahmer, N., Farese, R.V., Jr. & Walther, T.C. Balancing the fat: lipid droplets and human disease. EMBO Mol Med 5, 973–983 (2013).

4. Cruz, A.L.S., Barreto, E.A., Fazolini, N.P.B., Viola, J.P.B. & Bozza, P.T. Lipid droplets: platforms with multiple functions in cancer hallmarks. Cell Death Dis 11, 105 (2020).

5. Petan, T. Lipid Droplets in Cancer. Rev Physiol Biochem Pharmacol (2020).

6. Luo, W. et al. Adding fuel to the fire: The lipid droplet and its associated proteins in cancer progression. Int J Biol Sci 18, 6020–6034 (2022).

7. Heier, C. & Kuhnlein, R.P. Triacylglycerol Metabolism in Drosophila melanogaster. Genetics 210, 1163–1184 (2018).

8. Welte, M.A. & Gould, A.P. Lipid droplet functions beyond energy storage. Biochim Biophys Acta 1862, 1260–1272 (2017).

9. Gronke, S. et al. Brummer lipase is an evolutionary conserved fat storage regulator in Drosophila. Cell Metab 1, 323–330 (2005).

10. Kimmel, A.R. & Sztalryd, C. The Perilipins: Major Cytosolic Lipid Droplet-Associated Proteins and Their Roles in Cellular Lipid Storage, Mobilization, and Systemic Homeostasis. Annual review of nutrition 36, 471–509 (2016).

11. Itabe, H., Yamaguchi, T., Nimura, S. & Sasabe, N. Perilipins: a diversity of intracellular lipid droplet proteins. Lipids in health and disease 16, 83 (2017).

12. Islimye, E., Girard, V. & Gould, A.P. Functions of Stress-Induced Lipid Droplets in the Nervous System. Front Cell Dev Biol 10, 863907 (2022).

13. Van Den Brink, D.M., et al. Physiological and pathological roles of FATP-mediated lipid droplets in Drosophila and mice retina. PLoS Genet 14, e1007627 (2018).

14. Bailey, A.P. et al. Antioxidant Role for Lipid Droplets in a Stem Cell Niche of Drosophila. Cell 163, 340–353 (2015).

15. Ioannou, M.S. et al. Neuron-Astrocyte Metabolic Coupling Protects against Activity-Induced Fatty Acid Toxicity. Cell 177, 1522–1535.e1514 (2019).

16. Liu, L., MacKenzie, K.R., Putluri, N., Maletic-Savatic, M. & Bellen, H.J. The Glia-Neuron Lactate Shuttle and Elevated ROS Promote Lipid Synthesis in Neurons and Lipid Droplet Accumulation in Glia via APOE/D. Cell Metab 26, 719–737.e716 (2017).

17. Danielli, M., Perne, L., Jarc Jovicic, E. & Petan, T. Lipid droplets and polyunsaturated fatty acid trafficking: Balancing life and death. Front Cell Dev Biol 11, 1104725 (2023).

18. Boren, J. & Brindle, K.M. Apoptosis-induced mitochondrial dysfunction causes cytoplasmic lipid droplet formation. Cell Death Differ 19, 1561–1570 (2012).

19. Hakumaki, J.M. & Kauppinen, R.A. 1H NMR visible lipids in the life and death of cells. Trends Biochem Sci 25, 357–362 (2000).

20. Bischof, J. et al. Clearing the outer mitochondrial membrane from harmful proteins via lipid droplets. Cell death discovery 3, 17016 (2017).

21. Yacobi-Sharon, K., Namdar, Y. & Arama, E. Alternative Germ Cell Death Pathway in Drosophila Involves HtrA2/Omi, Lysosomes, and a Caspase-9 Counterpart. Dev Cell 25, 29–42 (2013).

22. Napoletano, F. et al. p53-dependent programmed necrosis controls germ cell homeostasis during spermatogenesis. PLoS Genet 13, e1007024 (2017).

23. Zohar-Fux, M. et al. The phagocytic cyst cells in Drosophila testis eliminate germ cell progenitors via phagoptosis. Sci Adv 8, eabm4937 (2022).

24. Chao, C.F. et al. An important role for triglyceride in regulating spermatogenesis. Elife 12 (2024).

25. Wang, C. & Youle, R.J. Predominant requirement of Bax for apoptosis in HCT116 cells is determined by Mcl-1’s inhibitory effect on Bak. Oncogene 31, 3177–3189 (2012).

26. Ozturk-Colak, A. et al. FlyBase: updates to the Drosophila genes and genomes database. Genetics 227 (2024).

27. Sztalryd, C. & Brasaemle, D.L. The perilipin family of lipid droplet proteins: Gatekeepers of intracellular lipolysis. Biochimica et biophysica acta. Molecular and cell biology of lipids 1862, 1221–1232 (2017).

28. Skinner, J.R. et al. Diacylglycerol enrichment of endoplasmic reticulum or lipid droplets recruits perilipin 3/TIP47 during lipid storage and mobilization. J Biol Chem 284, 30941–30948 (2009).

29. Miner, G.E. & Cohen, S. Protocol for monitoring fatty acid trafficking from lipid droplets to mitochondria in cultured cells. STAR Protoc 5, 103236 (2024).

30. Miura, S. et al. Functional conservation for lipid storage droplet association among Perilipin, ADRP, and TIP47 (PAT)-related proteins in mammals, Drosophila, and Dictyostelium. J Biol Chem 277, 32253–32257 (2002).

31. Beller, M. et al. PERILIPIN-dependent control of lipid droplet structure and fat storage in Drosophila. Cell Metab 12, 521–532 (2010).

32. Zhao, X. et al. An RDH-Plin2 axis modulates lipid droplet size by antagonizing Bmm lipase. EMBO Rep 23, e52669 (2022).

33. Leist, M., Single, B., Castoldi, A.F., Kuhnle, S. & Nicotera, P. Intracellular adenosine triphosphate (ATP) concentration: a switch in the decision between apoptosis and necrosis. J Exp Med 185, 1481–1486 (1997).

34. Sênos Demarco, R., Uyemura, B.S., D’Alterio, C. & Jones, D.L. Mitochondrial fusion regulates lipid homeostasis and stem cell maintenance in the Drosophila testis. Nat Cell Biol 21, 710–720 (2019).

35. Buszczak, M., Lu, X., Segraves, W.A., Chang, T.Y. & Cooley, L. Mutations in the midway gene disrupt a Drosophila acyl coenzyme A: diacylglycerol acyltransferase. Genetics 160, 1511–1518 (2002).

36. Bersuker, K. et al. A Proximity Labeling Strategy Provides Insights into the Composition and Dynamics of Lipid Droplet Proteomes. Dev Cell 44, 97–112.e117 (2018).

37. Lee, E.F. et al. Crystal structure of ABT-737 complexed with Bcl-xL: implications for selectivity of antagonists of the Bcl-2 family. Cell Death Differ 14, 1711–1713 (2007).

38. Xu, X., Farach-Carson, M.C. & Jia, X. Three-dimensional in vitro tumor models for cancer research and drug evaluation. Biotechnol Adv 32, 1256–1268 (2014).

39. Suzuki, M., Youle, R.J. & Tjandra, N. Structure of Bax: coregulation of dimer formation and intracellular localization. Cell 103, 645–654 (2000).

40. Westphal, D. et al. Apoptotic pore formation is associated with in-plane insertion of Bak or Bax central helices into the mitochondrial outer membrane. Proc Natl Acad Sci U S A 111, E4076–4085 (2014).

41. Reyna, D.E. et al. Direct Activation of BAX by BTSA1 Overcomes Apoptosis Resistance in Acute Myeloid Leukemia. Cancer Cell 32, 490–505 e410 (2017).

42. Henne, W.M. & Cohen, S. Heterogeneity, dynamics and organelle interactions of lipid droplets. Nat Rev Mol Cell Biol (2026).

43. Friday, C.M. et al. APOE4 reshapes the lipid droplet proteome and modulates microglial inflammatory responses. Neurobiol Dis 212, 106983 (2025).

44. Pereira-Dutra, F.S. & Bozza, P.T. Lipid droplets diversity and functions in inflammation and immune response. Expert Rev Proteomics 18, 809–825 (2021).

45. Kumar, J. et al. Lipid droplet-induced T cell death sustains autoimmune tissue inflammation. Cell Metab (2026).

46. Liu, L. et al. Glial Lipid Droplets and ROS Induced by Mitochondrial Defects Promote Neurodegeneration. Cell 160, 177–190 (2015).

47. Braun, T. et al. Apoptosis-resistant cells drive compensatory proliferation via cell-autonomous and non-autonomous functions of the initiator caspase Dronc. Nat Commun 16, 10871 (2025).

48. Rueden, C.T. et al. ImageJ2: ImageJ for the next generation of scientific image data. BMC Bioinformatics 18, 529 (2017).

49. Fanfone, D. et al. Confined migration promotes cancer metastasis through resistance to anoikis and increased invasiveness. Elife 11 (2022).

50. Thiam, A.R. et al. COPI buds 60-nm lipid droplets from reconstituted water-phospholipid-triacylglyceride interfaces, suggesting a tension clamp function. Proc Natl Acad Sci U S A 110, 13244–13249 (2013).

51. Chorlay, A. & Thiam, A.R. An Asymmetry in Monolayer Tension Regulates Lipid Droplet Budding Direction. Biophys J 114, 631–640 (2018).

52. Hughes, C.S. et al. Single-pot, solid-phase-enhanced sample preparation for proteomics experiments. Nat Protoc 14, 68–85 (2019).

53. Rouchidane Eyitayo, A., Daury, L., Priault, M. & Manon, S. The membrane insertion of the pro-apoptotic protein Bax is a Tom22-dependent multi-step process: a study in nanodiscs. Cell death discovery 10, 335 (2024).

54. Rambold, A.S., Cohen, S. & Lippincott-Schwartz, J. Fatty acid trafficking in starved cells: regulation by lipid droplet lipolysis, autophagy, and mitochondrial fusion dynamics. Dev Cell 32, 678–692 (2015).

