## Supplemental figure legends for "Lipid droplets sequester cell death effectors and delay regulated cell death execution"

**Fig. S1: BxPC3 and HCT116 cells competence to different cell death modalities**

**(A-E)** Kinetics of cell death of BxPC3 cells or **(F-I)** of HCT116 cells monitored by Sytox Green uptake. Graphs represent cell death (Sytox-positive cells normalized to confluency) in the function of time. BxPC3 cells were treated with the indicated compounds, and cell death was tracked in real time using the IncuCyte live-cell imaging system. Treatments included: intrinsic apoptosis induced by Actinomycin D (ActD, 1 μM) + ABT-737 (1 μM), extrinsic apoptosis induced by cycloheximide (CHX, 5 μg/mL) + tumor necrosis factor alpha (TNFα, 20 ng/mL), necrosis induced by ionomycin (25 μM), and necroptosis by the SMAC mimetics (BV6,10 μM) + TNFα + caspase inhibitor ZVAD (10 μM). Caspase inhibitor (QVD, 10 μM) was used to inhibit apoptosis, while necrosulfonamide (NSA, 5 μM) to prevent necroptosis.

**Fig. S2: LDs accumulate during physiological or caspase-induced germ cell death**

**(A)** A schematic diagram showing the apical tip of the *Drosophila* testis. Germ stem cells (in pink) are localized apically in the hub region. Goniablast (in yellow) divide into spermatogonia (in orange), which undergo four rounds of mitosis and occasionally spontaneous cell death (in red). Cyst stem cells and cyst cells are shown in dark and light blue, respectively.

**(B)** Representative images of the apical tip of testes from animals expressing nuclear-localized RFP (red) and Lamin-GFP (green) under early germ cell driver (*nos-Gal4*). Dying germ cells are observed in brightfield images. Nuclei are shown with DAPI staining (gray). Solid white squares indicate regions of interest, which are shown at higher magnification in the bottom panels. Dying germ cell cysts labeled with white arrows show the loss of Lamin-GFP and of nuclear localized RFP. Scale bars: 20 µm; magnified images: 10 µm.

**(C)** Representative images of testes from *vasa*-EGFP flies. Dying germ cells are observed in brightfield images. EGFP-positive germ cells are shown in green. LDs are stained with LipidTOX (red), and nuclei are shown with DAPI staining (gray). Solid white squares indicate regions of interest, which are shown at higher magnification in the bottom panels. Dying germ cell cyst is outlined with white dashed lines. Scale bars: 20 µm; magnified images: 10 µm.

**(D)** Representative images of testes dissected from animals expressing mCD8-GFP (green) under somatic cyst cell driver *C587-Gal4*. Dying germ cells are observed in brightfield images. GFP-positive cyst cells are shown in green. LDs are stained with LipidTOX (red), and nuclei are shown with DAPI staining (gray). Solid white squares indicate regions of interest, which are shown at higher magnification in the bottom panels. Scale bars: 20 µm; magnified images: 10 µm.

**(E)** Representative images of testes dissected from animals expressing either LacZ (control) or active form of Drice under early germ cell driver *nos-Gal4*. Dying germ cells are observed in brightfield images. LDs are stained with BODIPY$^{493/503}$ (green). Dying germ cell cysts are indicated with white arrows. Scale bars: 20 µm.

**(F)** Quantification of the number of dying germ cell cysts in the testes expressing either *LacZ* or active form of *Drice* under early germ cell driver *nos-Gal4*. Data represent the mean ± SD from three independent experiments (N = 3, n = 24 for each genotype). Statistical analysis was performed using two-tailed unpaired Student's t-test: ****, $p < 0.0001$.

**(G)** Quantification of average LD size in testes expressing either *LacZ* and active form of *Drice* under early germ cell driver *nos-Gal4*. Data represent the mean ± SD from three independent experiments (N = 3, each dot represents a single testis; n = 24 for each genotype). Statistical analysis was performed using two-tailed unpaired Student's t-test: **, $p < 0.01$.

**Fig. S3: Accumulating LDs labeled with perilipins in dying human cancer cell**

**(A)** *PLIN1-5* gene expression in BxPC3 and HCT116 cells. Data were normalized to *HPRT* reference gene.

**(B)** PLIN2 and **(C)** PLIN3 protein levels in control (DMSO) or cell death–induced BxPC3 cells. Protein lysates were collected 18 h after treatment. Treatments included: intrinsic apoptosis induced by Actinomycin D (ActD, 1 µM) + ABT-737 (1 µM), extrinsic apoptosis induced by cycloheximide (CHX, 5 µg/mL) + tumor necrosis factor alpha (TNFα, 20 ng/mL). Densitometric analysis of immunoblots is shown; data represent the mean ± SD from three independent experiments (N = 3). Statistical analysis was performed using one-way ANOVA, ns, no significance.

**(D)** Quantification of percentage of LDs (BODIPY$^{493/503}$) showing at least 20% overlap with PLIN3 (left) or PLIN2 (right) staining. Intrinsic apoptosis was induced by treatment with ActD + ABT-737 (1 µM each) in the presence of QVD in order to prevent cell detachment during staining.

**(E)** Representative western blot images for detection of ATGL in HCT116 and BxPC3 cells. Vinculin was used as a loading control.

**(F)** Representative LDs fractions analyzed by western blot. PLIN2 is enriched in fractions 1 to 4. Cox IV (cytochrome c oxidase IV), an inner mitochondrial membrane protein, is detected in the last fraction, while GAPDH (glyceraldehyde-3-phosphate dehydrogenase), a cytosolic protein, is detected from fraction 11. IP = input.

**(G)** Quantification of total LDs amount, measured by BODIPY$^{493/503}$ intensity, isolated from HCT116 cells following intrinsic apoptosis induction with ActD + ABT-737 (1 µM each) for 5 h. Only fractions 1 and 2 were used to evaluate BODIPY$^{493/503}$ intensity normalized to Vinculin. Data represent the mean ± SD from four independent experiments (N = 4). Statistical analysis was performed using two-tailed unpaired Student's t-test: *, $p < 0.05$.

**Fig. S4: Accumulating LDs require DGAT1/2 activity in dying human cancer cells**

**(A)** Representative fluorescence images of HCT116 cells treated with a combination of 15 µM DGAT1 and 10 µM DGAT2 inhibitors (DGATi) to block *de novo* LD formation. Cell death was induced in the presence or absence of DGATi for 18 h. LDs were visualized using BODIPY$^{93/503}$ or LipidTOX (green), and nuclei were stained with Hoechst 33342 (blue). Scale bars: 20 µm.

**(B)** Quantification of the mean LDs count per cell. Data represent the mean ± SD from three independent experiments (N = 3). Statistical analysis was performed using two-way ANOVA. ****, $p < 0.0001$.

**(C)** Total LD area per cell (µm2). Data represent the mean ± SD from three independent experiments (N = 3). Statistical analysis was performed using two-way ANOVA. ****, $p < 0.0001$.

**(D, E)** Western blot images showing PLIN2, PLIN3, and ATGL levels in the LD fraction following inhibition of DGAT1/2 activity during intrinsic apoptosis induction. HCT116 WT cells were treated either with DMSO (control) or Actinomycin D (ActD, 1 µM), ABT-737 (1 µM) and 10 µM of each DGAT1 and DGAT2 inhibitor (DGATi).

**(E)** Pooled sucrose gradient fractions excluding the LD fractions (1-4) for Vinculin (cell loading control) and PARP-1 cleavage detection.

**(F, G, H)** Quantification of PLIN3, PLIN2 and ATGL at the LDs. Densitometry analysis was performed on fractions 1 and 2. Statistical analysis was performed using two-tailed unpaired Student's t-test: *, $p < 0.05$.

**Fig. S5: dPlin1, Bmm labeling and *mdy* requirement in LDs of *Drosophila* germ cells**

**(A)** Representative images of the apical tip of testes from 5-day-old $w^{1118}$ flies. Dying germ cells are observed in brightfield images. LDs were stained with BODIPY$^{493/503}$ (green) and dPlin1 using antibody staining (red). Solid white squares indicate regions of interest, which are shown at higher magnification in the bottom panels. Scale bars: 20 μm; magnified images: 10 μm.

**(B)** Representative images of fat body from 5-day-old Bmm-GFP flies. Endogenous Bmm-GFP proteins are shown in green on LDs (LipidTOX staining, magenta). Scale bars: 20 μm.

**(C)** Representative images of the apical tip of testes from Bmm-GFP flies. LDs were stained with BODIPY$^{493/503}$ (green) and Bmm-GFP using antibody against GFP (red). Solid white squares (ROI 1) indicate regions of healthy cells, which are shown at higher magnification in the middle panels. Solid white squares (ROI 2) indicate regions of dying cells, which are shown at higher magnification in the bottom panels. Bmm-positive LDs are shown with arrows. Scale bars: 20 μm; magnified images: 10 μm.

**(D)** Representative images of testes from 5-day-old *wild-type* ($w^{1118}$) and *midway* mutant ($mdy^{QX25}$) flies. LDs were stained with BODIPY$^{93/503}$ (green). Scale bars: 20 μm.

**(E)** Quantification of average LD size in the testes. Data represent the mean ± SD from three independent experiments (N = 3, each dot represents a single testis; n = 15 and 13 for each genotype, respectively). Statistical analysis was performed using two-tailed unpaired Student's t-test: ****, $p < 0.0001$.

**Fig S6: *CPT2* knockdown and etomoxir does not affect cell death in *Drosophila* germ cells and cancer cells, respectively**

**(A)** Representative images of testes dissected from 3-day-old animals expressing either GFP RNAi (control) or CPT2 RNAi under germ cell driver Bam-Gal4. The testes were stained with DAPI (green), Traffic jam (Tj, red) and Fas III (magenta). Scale bars: 20 µm.

**(B)** Graph showing germ stem cell (GSC) number. Data represent the mean ± SD from three independent experiments (N = 3, each dot represents a single testis; n = 26 and 25 for each genotype). Statistical analysis was performed using two-tailed unpaired Student's t-test: ns, no significance.

**(C)** Quantification of the number of dying germ cell cysts in the testes expressing either *GFP RNAi* or *CPT2 RNAi* under germ cell driver *Bam-Gal4*. Data represent the mean ± SD from three independent experiments (N = 3, n = 26 for each genotype). Statistical analysis was performed using two-tailed unpaired Student's t-test: ns, no significance

**(D and E)** Graph representing the percentage of cell death (Sytox-positive cells normalized to confluency) of BxPC3 cells (D) and HCT116 (E) 20 h after cell death induction. Cell death was induced by ActD/ABT-737 treatment (1 µM of each) in the presence or absence of Etomoxir (100 µM). Two-way ANOVA statistical test was performed: ns, no significance.

**Fig S7: *mdy*$^{QX25}$ does not affect germ cell death in *Drosophila* testes**

**(A)** Representative images of the apical tip of testes from 5-day-old $w^{1118}$ and *mdy* mutant (*mdy*$^{QX25}$) flies. Dying germ cells are observed in brightfield images. Scale bars: 20 μm.

**(B)** Quantification of germ cell cysts in the testes. Data represent the mean ± SD from three independent experiments (N = 3, each dot represents a single testis; n = 39 for each genotype). Statistical analysis was performed using two-tailed unpaired Student's t-test: ns, no significance.

**Fig. S8: Bax localization at LDs and mitochondria during apoptosis**

**(A)** LD isolation from HCT116 cells with doxycycline (dox)-induced overexpression of Bax (Bax OE). Cells with overexpression of empty vector (EV) were used as a control. LDs were isolated as previously described from cells 5 h after induction of Bax overexpression via dox treatment (0.1 µg/µL). LD fractions were analyzed by western blot to detect Bax, Bcl-xL and PLIN3. The blot shows endogenous Bax and Flag-Bax with the lower and higher molecular weights, respectively. The cytosolic fraction was used for detection of Vinculin (loading control). This experiment is representative of two biological repeats (N = 2).

**(B)** Representative western blot analysis of cleaved PARP and Bax following doxycycline-induced Bax overexpression. This experiment is representative of two biological repeats (N = 2).

**(C)** Total cell lysates from attached cells (AC), floating cells (FC) and DMSO-treated control were analyzed by western blot to assess caspase-3 cleavage following treatment with ActD/ABT-737 (1 µM each) for 5 h. This experiment is representative of three biological repeats (N = 3).

**(D)** Crude mitochondria were isolated from AC, FC and DMSO-treated control following treatment with ActD/ABT-737 (1 µM each) for 5 h. The mitochondria fractions were analyzed by western blot to detect Bax, Bcl-xL, cytochrome c (cyt c) and Cox IV (mitochondria loading control).

**(E and F)** Quantification of Bax and Bcl-xL levels in the mitochondrial fraction shown in (D). This experiment is representative of three biological repeats (N = 3). Statistical analysis was performed using two-tailed unpaired Student's t-test: *, $p < 0.05$; **, $p < 0.01$.

**(G)** Representative images of HCT116 cells overexpressing EGFP-Bax treated with ActD/ABT-737 (1 µM of each). Mitochondria were labeled with MitoTracker (magenta), and EGFP-Bax is shown in green. The refractive index channel was used to visualize LDs by holotomographic imaging. Scale bars: 5 µm.

**Fig. S9: Bax activation by BTSA1 promotes Bax recruitment to artificial lipid droplets (aLDs)**

**(A)** Representative images of artificial LDs (aLDs). Neutral lipid cores were stained with BODIPY$^{493/503}$ (green), while the surrounding phospholipids monolayer was labeled with Rhodamine phosphatidyl ethanolamine (red). Scale bar: 20μm.

**(B)** Representative western blot images showing Bax level on aLDs following Bax activation by BTSA1. HCT116 cytosol depleted of endogenous LDs was incubated with BODIPY$^{493/503}$-labeled aLDs in the presence of either 1 μM BTSA1 or DMSO. Following this incubation, aLDs were isolated, and the aLD-containing fractions (fractions 1–4) were collected for Western blot analysis. The remaining gradient fractions were pooled and used for vinculin detection as a cytosolic loading control (lysate).

**(C, D and E)** Quantification of Bax, Bcl-xL and PLIN3 at the aLDs. Only fractions 1 and 2 were used for densitometric analysis. aLD quantity in those fractions was evaluated by measuring BODIPY$^{493/503}$ intensity. Data represent the mean ± SD relative to BODIPY$^{493/503}$ intensity from three independent experiments (N = 3). Statistical analysis was performed using two-tailed unpaired Student's t-test: *, $p < 0.05$.
