## Supplementary figures and images for "Lipid droplets sequester cell death effectors and delay regulated cell death execution"

### Supplemental Figure 1

#### Supplementary Figure 1

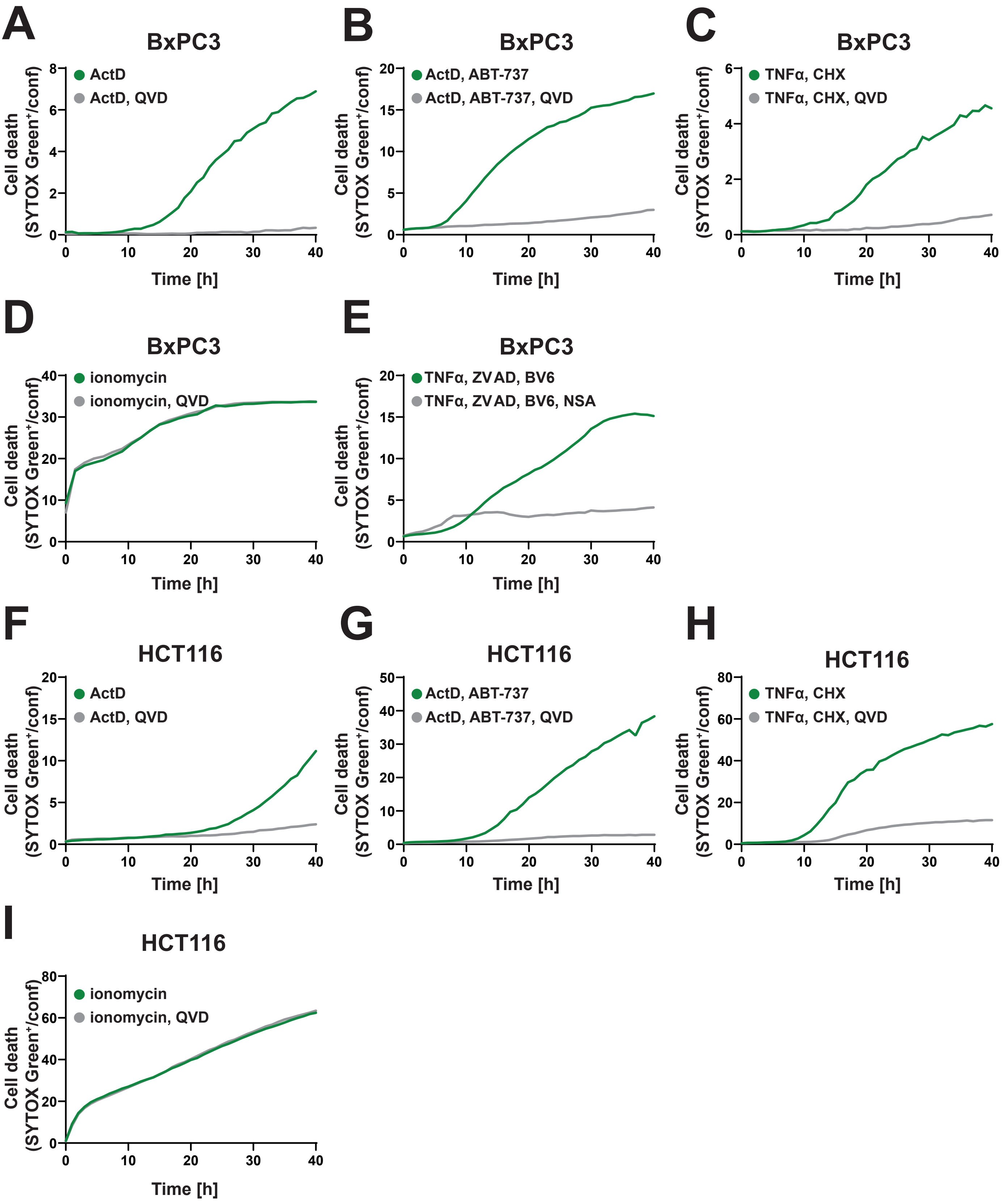

### Supplemental Figure 2

#### Supplementary Figure 2

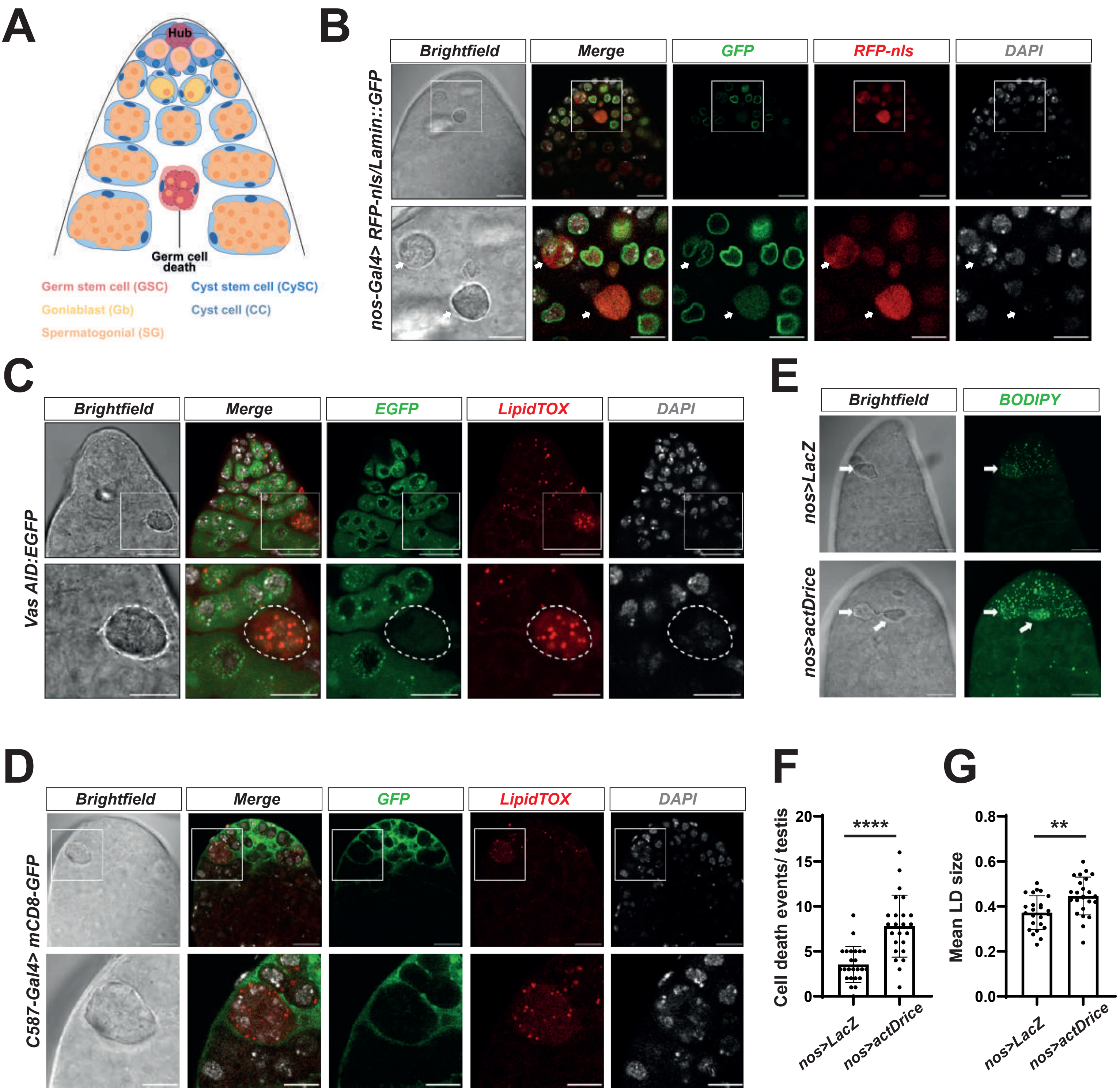

### Supplemental Figure 3

#### Supplementary Figure 3

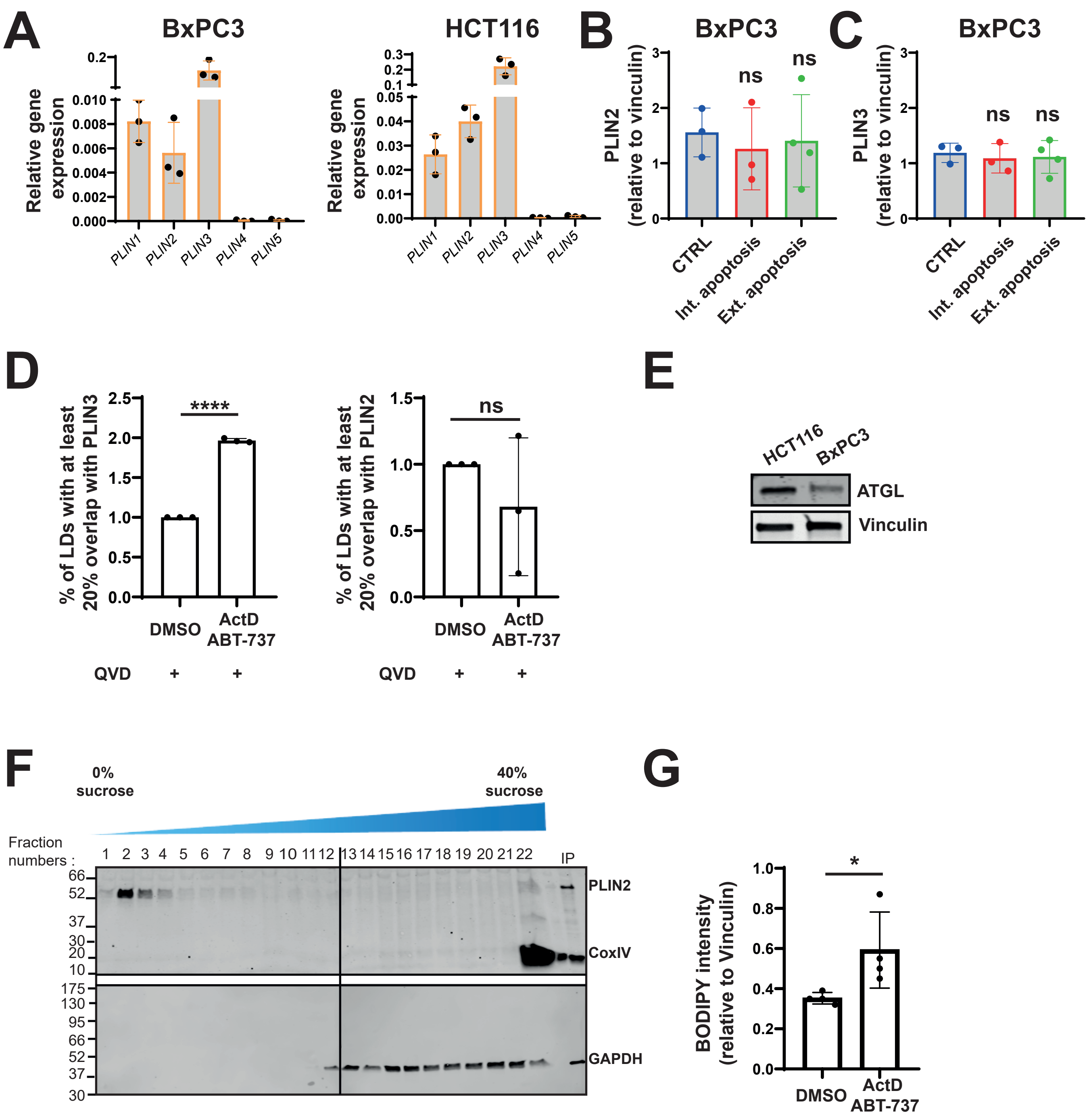

### Supplemental Figure 4

#### Supplementary Figure 4

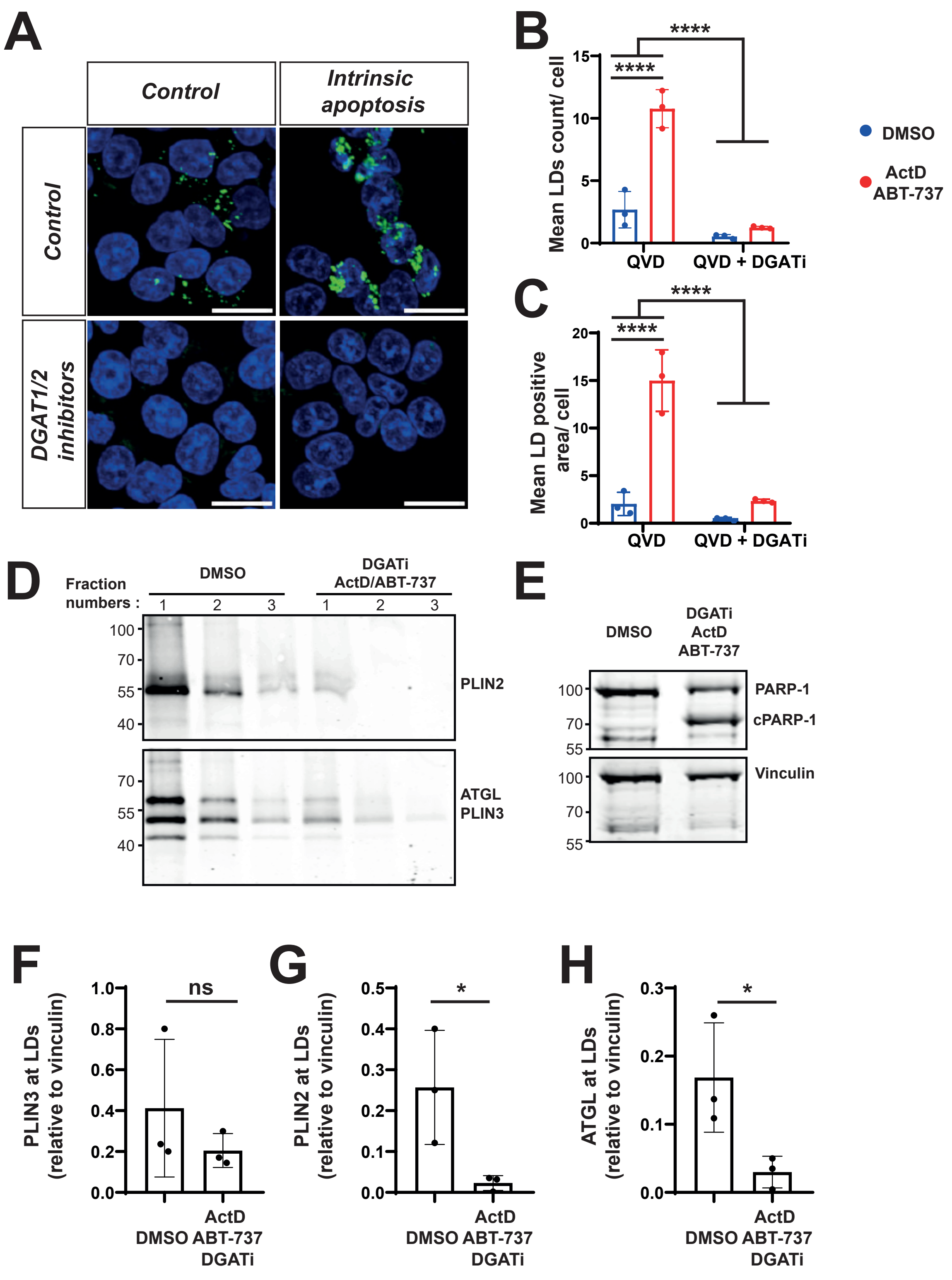

### Supplemental Figure 5

#### Supplementary Figure 5

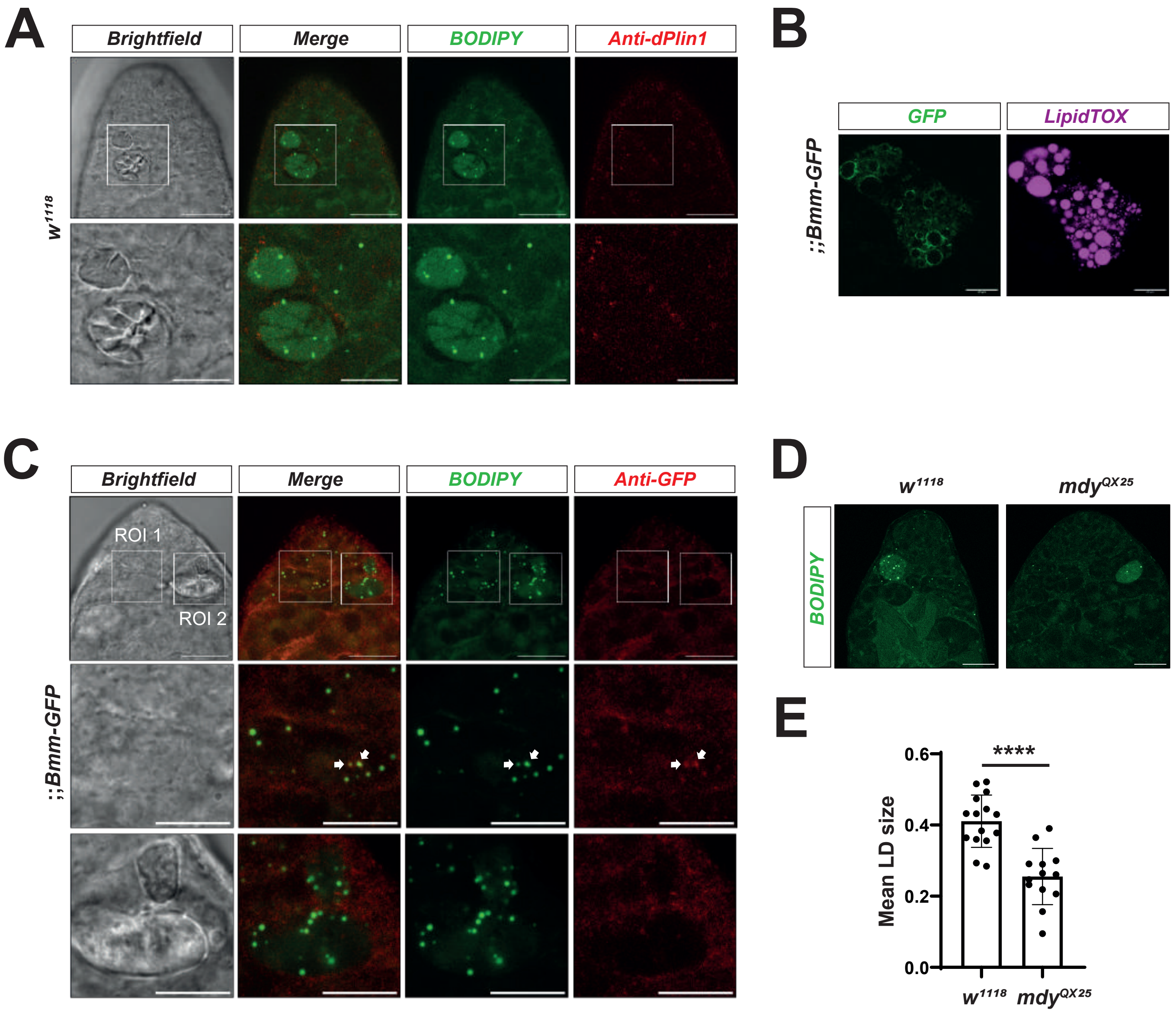

### Supplemental Figure 6

#### Supplementary Figure 6

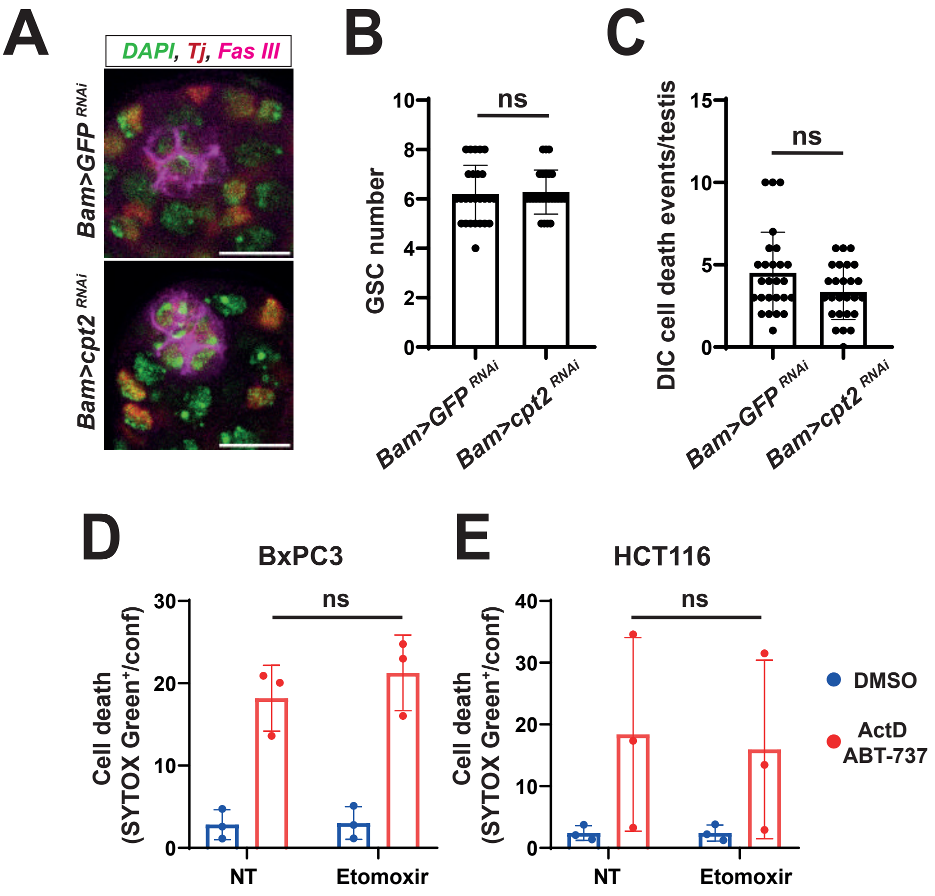

### Supplemental Figure 7

#### Supplementary Figure 7

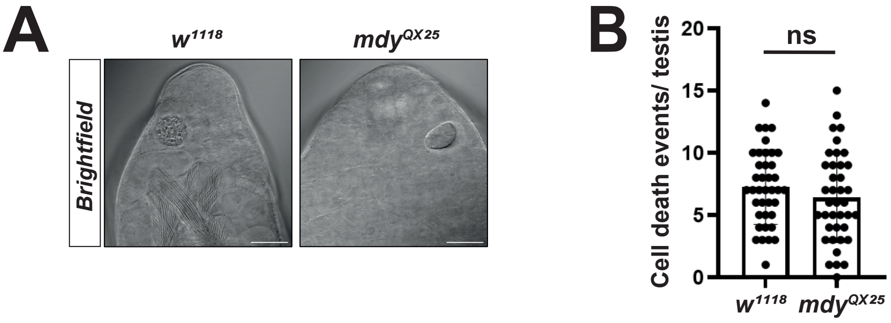

### Supplemental Figure 8

#### Supplementary Figure 8

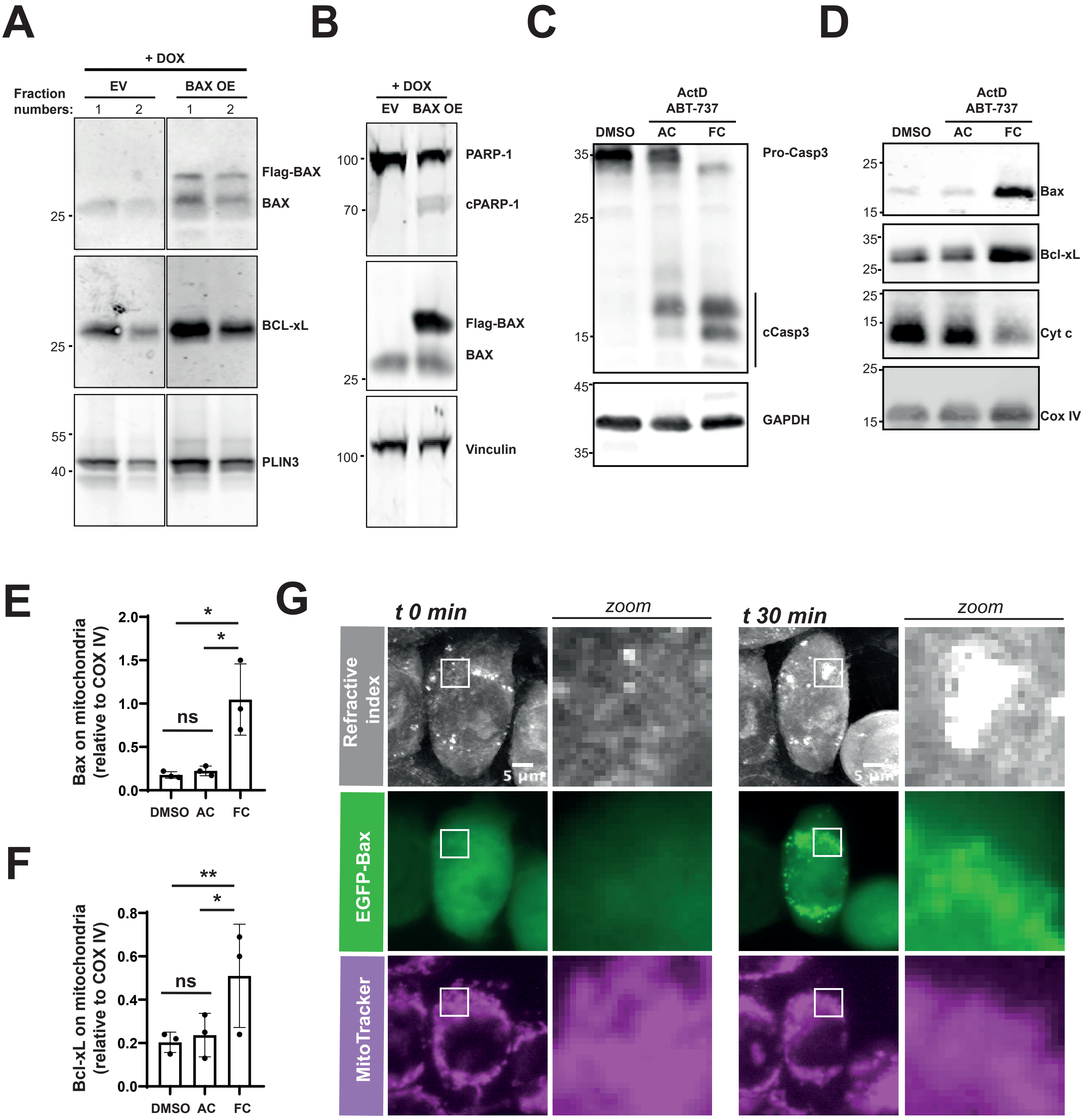

### Supplemental Figure 9

#### Supplementary Figure 9

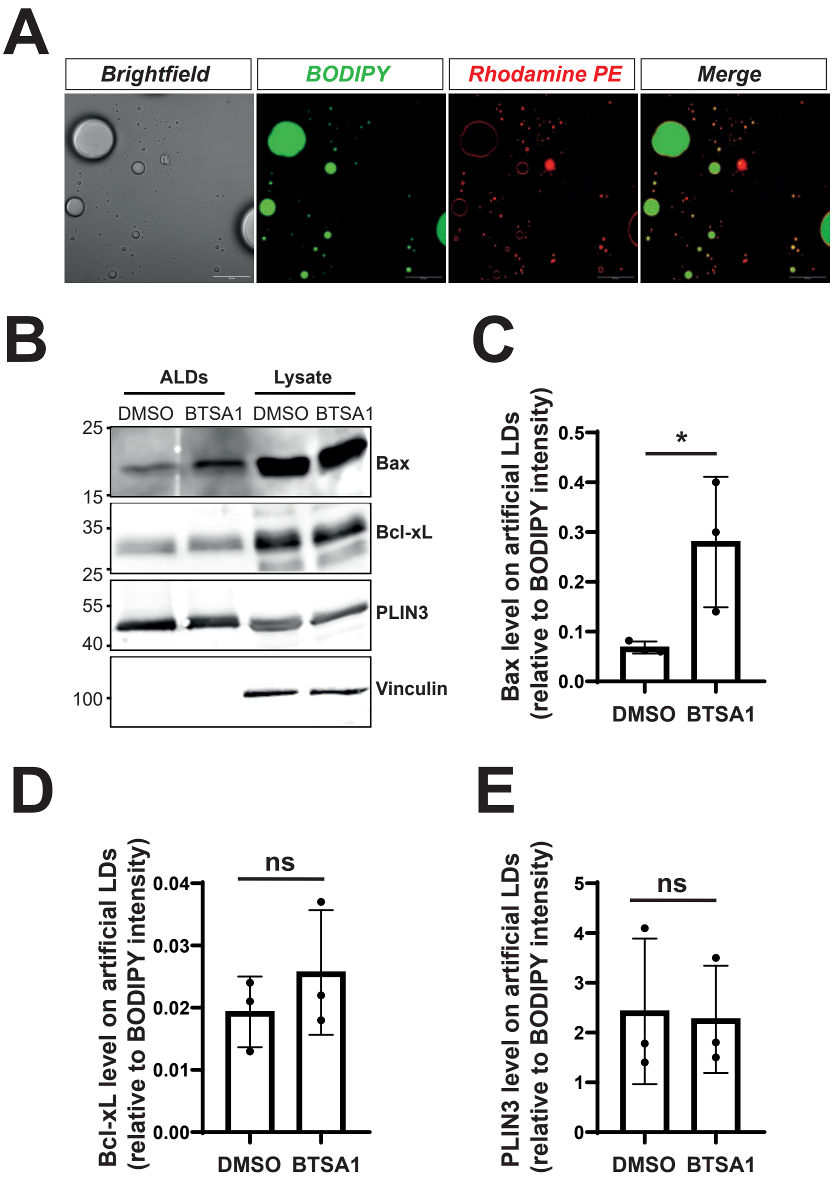
